# Npas1^+^-Nkx2.1^+^ Neurons Are an Integral Part of the Cortico-pallido-cortical Loop

**DOI:** 10.1101/644674

**Authors:** Zachary A. Abecassis, Brianna L. Berceau, Phyo H. Win, Daniela Garcia, Harry S. Xenias, Qiaoling Cui, Arin Pamucku, Suraj Cherian, Vivian M. Hernández, Uree Chon, Byung Kook Lim, Nicholas J. Justice, Raj Awatramani, Yongsoo Kim, Bryan M. Hooks, Charles R. Gerfen, Simina M. Boca, C. Savio Chan

## Abstract

Within the basal ganglia circuit, the external globus pallidus (GPe) is critically involved in motor control. Aside from Foxp2^+^ neurons and ChAT^+^ neurons that have been established as unique neuron types, there is little consensus on the classification of GPe neurons. Properties of the remaining neuron types are poorly-defined. In this study, we leverage new mouse lines, viral tools, and molecular markers to better define GPe neuron subtypes. We found that Sox6 represents a novel, defining marker for GPe neuron subtypes. Lhx6^+^ neurons that lack the expression of Sox6 were devoid of both parvalbumin and Npas1. This result confirms previous assertions of the existence of a unique Lhx6^+^ population. Neurons that arise from the Dbx1^+^ lineage were similarly abundant in the GPe and displayed a heterogeneous makeup. Importantly, tracing experiments revealed that Npas1^+^-Nkx2.1^+^ neurons represent the principal non-cholinergic, cortically-projecting neurons. In other words, they form the pallido-cortical arm of the cortico-pallido-cortical loop. Our data further described that pyramidal-tract neurons in the cortex collateralized within the GPe, forming a closed-loop system between the two brain structures. Overall, our findings reconcile some of the discrepancies that arose from differences in techniques or the reliance on pre-existing tools. While spatial distribution and electrophysiological properties of GPe neurons reaffirm the diversification of GPe subtypes, statistical analyses strongly support the notion that these neuron subtypes can be categorized under the two principal neuron classes—i.e., PV^+^ neurons and Npas1^+^ neurons.

**Significance statement:** The poor understanding of the neuronal composition in the GPe undermines our ability to interrogate its precise behavioral and disease involvements. In this study, twelve different genetic crosses were used, hundreds of neurons were electrophysiologically-characterized, and over 100,000 neurons were histologically- and/or anatomically-profiled. Our current study further establishes the segregation of GPe neuron classes and illustrates the complexity of GPe neurons in adult mice. Our results support the idea that Npas1^+^-Nkx2.1^+^ neurons are a distinct GPe neuron subclass. By providing a detailed analysis of the organization of the cortico-pallidal-cortical projection, our findings establish the cellular and circuit substrates that can be important for motor function and dysfunction.

## Introduction

The basal ganglia are a network of brain nuclei that are involved in motor control and adaptive behavior. Dysfunction within this circuit can be devastating, as seen in patients afflicted with Parkinson’s Disease (PD), Huntington’s Disease, and dystonias (Cox and Witten, 2019; DeLong and Wichmann, 2007; Dudman and Krakauer, 2016; Graybiel, 2008; Grillner and Robertson, 2016; Ito and Doya, 2011; Jahanshahi et al., 2015; Klaus et al., 2019; Mink, 2018; Nambu and Tachibana, 2014; Pennartz et al., 2009; Redgrave et al., 2010). The external globus pallidus (GPe) is a key nucleus within the basal ganglia. Decorrelated, phasic changes in GPe neuron activity are observed with normal movements (Anderson and Horak, 1985; Dodson et al., 2015; Jin et al., 2014; Mallet et al., 2016; Shi et al., 2004; Turner and Anderson, 2005). Alterations in the firing pattern of these neurons are associated with hypokinetic motor symptoms in both animal models of PD and human patients (Boraud et al., 1998; Chan et al., 2011; Filion et al., 1991; Hutchison et al., 1994; Jaeger and Kita, 2011; Magill et al., 2001; Mallet et al., 2008; Nini et al., 1995; Raz et al., 2000; Rothblat and Schneider, 1995).

Prior studies in the field have suggested GPe neuron subtypes are involved in some aspects of movement control (Dodson et al., 2015; Glajch et al., 2016; Mastro et al., 2017). However, precisely how these neuron subclasses are involved in motor function and dysfunction remains poorly-defined. Our past work characterized two principal classes of GPe neurons, parvalbumin-expressing (PV^+^) neurons and Npas1-expressing (Npas1^+^) neurons, which account for 55% and 27% of the GPe neurons, respectively. PV^+^ neurons project primarily to the subthalamic nucleus (STN) and Npas1^+^ neurons target the dorsal striatum (dStr). The Npas1^+^ population can be further broken down into distinct Foxp2-expressing (Foxp2^+^) and Nkx2.1-expressing (Nkx2.1^+^) subpopulations, with Foxp2^+^ neurons representing a unique population referred to as “arkypallidal” neurons (Abdi et al., 2015; Dodson et al., 2015; Hernandez et al., 2015). GPe neurons lacking Foxp2 expression, commonly referred to as “prototypic” neurons, are a more heterogeneous population. As we lack a complete description of the molecular identity of prototypic neurons, the precise function of prototypic neurons has not been systematically studied.

Lhx6-expressing (Lhx6^+^) neurons represent a substantial fraction of the prototypic GPe neuron subtype, although their reported abundance varies widely across laboratories (Abrahao and Lovinger, 2018; Dodson et al., 2015; Hernandez et al., 2015; Mastro et al., 2014). Unfortunately, due to limitations in the availability of a reliable transgenic mouse and antibodies to study this subset, a discrepancy remains in its abundance and extent of overlap with PV^+^ neurons and Npas1^+^ neurons across laboratories (Hegeman et al., 2016). In this study, we hypothesize the existence of a unique Lhx6^+^ GPe population that corresponds to the PV^−^ and Npas1^−^ (PV^−^-Npas1^−^) neurons we previously identified and accounts for ∼15–20% of the total GPe neuron population (Hernandez et al., 2015). These neurons could play an important role in neural function if they target a unique brain area, such as the cortex, which has been described (Ahrlund-Richter et al., 2019; Chen et al., 2015; Saunders et al., 2015; Schwarz et al., 2015; Sun et al., 2019; Van der Kooy and Kolb, 1985). With the advent of additional transgenic lines and viruses, we used molecular marker expression and connectome analysis to reconcile discrepancies and provide a more in-depth analysis of the neuronal makeup and its diversity within the GPe. We confirmed the existence of a unique Lhx6^+^ neuron population by its lack of Sox6 expression. This Lhx6^+^ population does not correspond to neurons that arise from the Dbx1-lineage, which is known to colonize the GPe. We found that Npas1^+^-Nkx2.1^+^ neurons represent the principal, non-cholinergic, cortically-projecting population, and they are part of a closed-loop formed between cortex and the GPe. We propose that Npas1^+^-Nkx2.1^+^ neurons, along with the previously identified Npas1^+^-Foxp2^+^ and ChAT^+^ neurons, are unique GPe neuron types.

## Methods

### Mice

All procedures were done in accordance with protocols approved by Northwestern University, the University of California at San Diego, University of Pittsburgh, and Janelia Research Campus Institutional Animal Care and Use Committees and were in compliance with the National Institutes of Health Guide to the Care and Use of Laboratory Animals. Experiments were conducted with the following mouse lines: LSL(Lox-STOP-Lox)-tdTomato (Ai14, Jax 007914), FSF(Frt-STOP-Frt)-LSL-tdTomato (Ai65, Jax 021875). Dbx1-Cre (Dbx1-ires-Cre, MMRRC 031751) (Harris et al., 2014) were crossed with LSL-tdTomato. As the Dbx1-Cre line is prone to germline recombination, recombination patterns were routinely monitored and compared against data on Allen’s Transgenic Characterization data portal (http://connectivity.brain-map.org/transgenic/experiment/100141723). Any mice displaying ectopic expression were excluded from subsequent analysis. Emx1-Cre (Emx1-ires-Cre, Jax 005628), Foxp2-Cre (Foxp2-ires-Cre, Jax 030541), Lhx6-GFP (Lhx6-GFP BAC, MMRRC 000246), Nkx2.1-Flp (Nkx2.1-ires-Flp, Jax 028577), Npas1-Cre-tdTom (Npas1-Cre-tdTomato BAC, Jax 027718), PV-Cre (PV-ires-Cre, Jax 017320), PV-Flp (PV-2A-Flp, Jax 022730), PV-tdTom (PV-tdTomato BAC, Jax 027395), Sim1-Cre (Sim1-Cre BAC, MMRRC 037650) and Tlx3-Cre (Tlx3-Cre BAC, MMRRC 036670) were all used in this study. FSF-tdTomato (Ai65F) was generated as previously described (Daigle et al., 2018; Yetman et al., 2019). In brief, FSF-LSL-tdTomato was crossed with EIIa-Cre (Jax 003724) to delete the LSL cassette. Sox6-Cre was generated by performing a germline deletion of the FSF cassette from our previously reported Sox6-FSF-Cre (Poulin et al., 2018), using CAG-Flp (Kanki et al., 2006) line. Dbx1;Ai14 referred to as Dbx1-L-tdTom. Nkx2.1-Flp;Ai65 referred to as Nkx2.1-F-tdTom. PV-Cre;Ai14 referred to as PV-L-tdTom. PV-Flp;Ai65F referred to as PV-F-tdTom. PV-Flp;Dbx1-Cre;Ai65 referred to as PV-Dbx1-FL-tdTom. Mice were backcrossed and only heterozygous and hemizygous mice were used throughout the study to minimize the potential alteration of the phenotypes in mice carrying the transgene alleles (Chan et al., 2012). Mice were group-housed in a 12 h light-dark cycle. Food and water were provided *ad libitum*. All mice were maintained by backcrossing with C57BL/6J breeders (Jax 000664). The genotypes of all transgenic mice were determined by tail biopsy followed by PCR to identify the presence of the relevant transgenes. Both male and female mice were used in this study.

### Stereotaxic injections

Standard injection procedures were used as described previously (Cui et al., 2016). In brief, mice at postnatal days 28–35 and 45–55 were used for viral tracing and retrograde tracing experiments, respectively, were anesthetized with isoflurane, and were immobilized on a stereotaxic frame (David Kopf Instruments). A small craniotomy (∼1 mm diameter) was made with a dental drill (Osada) for injection into the target (see Table 1) using a calibrated glass micropipette (VWR) at a rate of 0.3–0.5 μl/min. The micropipette was left *in situ* for 5–10 min postinjection to maximize viral retention and to decrease capillary spread upon pipette withdrawal. The following adeno-associated viruses (AAVs) were used in this study: AAV-EF1a-CreOn-hChR2(H134R)-EYFP (Addgene viral prep #20298-AAV9) and AAV-hSyn-CreOn-mCherry (Addgene viral prep #50459-AAV8) were used to infect GPe neurons. AAVretro-ChR2-eYFP (Addgene viral prep #20298-AAVrg) was used for retrograde delivery of ChR2 in Emx1-Cre mice. To examine cortical neuron subtype-specific projections, Tlx3-Cre and Sim1-Cre mice were injected at around postnatal day 37. 30 nL AAV-flex-XFPs was injected per site. Mouse brains were then fixed by transcardial perfusion 2–3 weeks post-injection (Hooks et al., 2018). Mice injected with Alexa-conjugated cholera-toxin B subunit (CTb; Thermo Fisher Scientific), lentivirus (LVretro-Cre) (Knowland et al., 2017), or AAVs were processed for immunohistological analysis (see below) 7–14 d and 28–42 d after injection, respectively. For LV tracing experiments, CTb was injected in conjunction with LV to visualize targeting accuracy. Mice with injection outside of the targeted area were excluded from subsequent analysis.

**Table 1.** Injection coordinates

|  | Injection volume (nl) | P28–35 |  |  | P45–55 |  |  |
| --- | --- | --- | --- | --- | --- | --- | --- |
|  |  | AP (mm) | ML (mm) | DV (mm) | AP (mm) | ML (mm) | DV (mm) |
| Primary somatosensory, primary somatomotor (MOp, SSp) | 180 | +2.60 | ±2.16 | –2.55, –2.75 | +2.60 | ±2.16 | –2.25, –2.50 |
| Secondary somatomotor (MOs), rostral | 180 | +1.80 | ±0.96 | –1.20, –1.60 | +2.60 | ±2.04 | –2.55, –2.75 |
| Secondary somatomotor (MOs), caudal | – | – | – | – | +1.80 | ±0.96 | –1.20, –1.60 |
| Anterior cingulate cortex (AC) | 90 | – | – | – | +1.00 | ±0.40 | –1.60 |
| Agranular cortex (AG) | 180 | +2.25 | ±1.92 | –2.25, –3.00 | –2.60 | ±2.04 | –2.55, –2.75 |
| Dorsal striatum (dStr) | 180 | +0.70 | ±2.30 | –3.00, –3.40 | +0.70 | ±2.30 | –3.00, –3.40 |
| External globus pallidus (GPe) | 90 | –0.28 | ±2.15 | –4.10 | –0.28 | ±2.15 | –4.10 |
| Subthalamic nucleus (STN) | 90 | –1.45 | ±1.70 | –4.52 | –1.45 | ±1.70 | –4.53 |
| Substantia nigra (SN) | 180 | –2.65, –3.00 | ±1.50 | –4.50 | –2.95 | ±1.44 | –4.50 |
| Parafascicular nucleus (Pf) | 90 | –2.10 | ±0.64 | –3.50 | –2.15 | ±0.64 | –3.50 |

Coordinates are in relation to bregma. Negative AP values refer to coordinates caudal to bregma. Abbreviations: AP = anterior-posterior, ML = medial-lateral, DV = dorsal-ventral.

### Immunolabeling and quantification

Mice ranging in age from postnatal day 55–80 were anesthetized deeply with a ketamine-xylazine mixture and perfused transcardially first with phosphate-buffered saline (PBS) followed by fixative containing 4% paraformaldehyde (PFA, pH 7.4). Tissue was then postfixed in the same fixative for 2 h at 4 °C. Tissue blocks containing the GPe were sectioned using a vibrating microtome (Leica Instruments) at a thickness of 60 μm. Floating sections were blocked with 10% (v/v) normal goat or donkey serum (Thermo Fisher Scientific) and 0.2% (v/v) Triton X-100 in PBS for 30–60 min, and were subsequently incubated with primary antibodies (see Table 2) in the same solution for 16–24 h at 4 °C. After washes in PBS, the sections were incubated with Alexa-conjugated, secondary antibodies (Thermo Fisher Scientific, 1:500 dilution) at room temperature for 2 h. The sections were then washed, mounted with ProLong Antifade mounting medium (Thermo Fisher Scientific), and coverslipped. In a subset of the experiments, DAPI was used to delineate cytoarchitecture of different brain structures. Fluorescent images of injection sites were captured on an epifluorescence microscope (Keyence Corporation) using a 2x or 10× 0.45 numerical aperture (NA) objective. Immunoreactivity in neurons was examined on a laser-scanning confocal microscope (Olympus). For cell quantification, images of the entire GPe were acquired on a laser-scanning confocal microscope with a 60× 1.35 NA oil-immersion objective. Images encompassing the GPe were taken and stitched using FLUOVIEW Viewer (Olympus) or Photoshop (Adobe Systems). Cell counting was performed manually using the cell-counter plugin within Fiji (Schindelin et al., 2012). Cell counts were obtained from ∼28 optical sections that were captured at 1 µm increments (median = 28). Neurons were defined by cells that were immuno-positive for HuCD or NeuN (Hernandez et al., 2015). GPe sections from three different equally-spaced (400 µm) lateromedial levels (∼2.5, 2.1, and 1.7 mm from bregma) were sampled and assigned as lateral, intermediate, and medial, respectively (Hernandez et al., 2015). They correspond approximately to sagittal plate 7, 9, and 11 on the Allen reference atlas (http://mouse.brain-map.org/static/atlas). In this study, the GPe is considered to be the structure that spans between the dorsal striatum and the internal capsule, which define the rostral and caudal and borders of the GPe on a sagittal plane, respectively. The cytoarchitecture of the ventral border is more ambiguous. For consistency, six non-overlapping z-stacks (212.13 x 212.13 µm) traversing the long axis of the GPe were used to capture its dorsoventral extent. This strategy coincides with the ventral border demarcated by the dense astrocytic labeling in the Dbx1-L-tdTom mice (see Results) and that defined in the common reference atlas.

**Table 2.** Primary antibodies used in this study

| Antigen | Host species | Clonality | Source | Catalog no. | Lot. no. | Dilution | Working concentration |
| --- | --- | --- | --- | --- | --- | --- | --- |
| ChAT | Gt | Polyclonal | Millipore-Sigma | AB144P | 2916187 | 1:1,000 |  |
| ChAT | Rb | Polyclonal | Synaptic Systems | 297013 | 297013/3 | 1:2,000 | 0.5 µg/ml |
| Cre | Rb | Polyclonal | Synaptic Systems | 257003 | 257003/1-7 | 1:500 | 2.0 µg/ml |
| CTb | Gt | Polyclonal | List Biological Labs | 102946-502 | 7032A10 | 1:5,000-1:10,000 |  |
| DsRed | Rb | Polyclonal | Clontech | 632496 |  | 1:500-1:1,000 |  |
| Foxp2 | Ms | Monoclonal | Millipore-Sigma | MABE415 | Q2273099 | 1:500 | 1.0 µg/ml |
| Foxp2 | Rb | Monoclonal | Millipore-Sigma | ABE73 | 3037985 | 1:500 |  |
| Foxp2 | Rb | Polyclonal | Millipore-Sigma | HPA000382 | B115858 | 1:500 |  |
| GFP | Ck | Polyclonal | Abcam | ab13970 | GR3190550-9 | 1:1,000 |  |
| HuC/D | Ms | Monoclonal | Life Technologies | A21271 | 1900217 | 1:1,000 | 0.2 µg/ml |
| mCherry | Rt | Monoclonal | Life Technologies | M11217 | S1259077 | 1:1,000-1:5,000 | 0.4-2.0 µg/ml |
| NeuN | Rb | Polyclonal | Biosensis | R-3770-100 | R-3770-300-201605-SH | 1:1,000 | 1.0 ng/µl |
| Nkx2.1 | Rb | Polyclonal | Millipore-Sigma | 07-601 | 2887266 | 1:500 |  |
| Npas1* | Gp | Polyclonal |  |  |  | 1:5,000 |  |
| Npas1* | Rb | Polyclonal |  |  |  | 1:5,000 |  |
| Parvalbumin | Ms | Monoclonal | Millipore-Sigma | P3088 | 016M-4847V | 1:500 | 10.0 µg/ml |
| Parvalbumin | Ck | Polyclonal | Synaptic Systems | 195006 | 195006/2 | 1:2,000 |  |
| Parvalbumin | Gp | Polyclonal | Synaptic Systems | 195004 | 195004/1-19 | 1:1,000-1:2,000 |  |
| Parvalbumin | Rb | Polyclonal | Swant | PV 27 | 2014 | 1:1,000 |  |
| RFP | Ms | Monoclonal | Rockland | 200-301-379 | 34537 | 1:1,000 | 0.5 µg/ml |
| Sox6 | Rb | Polyclonal | Abcam | ab30455 | GR289554-2 | 1:5,000 | 0.2 µg/ml |
| tdTomato | Rt | Monoclonal | Kerafast | EST203 | 091918 | 1:1,000 | 0.5 µg/ml |
| VGLuT1 | Gp | Polyclonal | Millipore-Sigma | AB5905 |  | 1:1,000 |  |

\*See Hernandez et al. 2015. Abbreviations: Ck = chicken, Gp = guinea pig, Gt = goat, Ms = mouse, Rb = rabbit, Rt = rat.

To mathematically represent the spatial distribution of GPe neurons and to compare that across neuron populations, sagittal brain sections were histologically processed. Images were manually aligned to the Allen Reference Atlas based on the structural information across the entire brain section. Images were transformed linearly (i.e. rotation, scaling) with no warping algorithms applied. Neurons that were not present within the confines of the GPe in the reference atlas were removed from subsequent analysis. GPe neurons located at lateral, intermediate, and medial levels (∼2.5, 2.1, and 1.7 mm lateral from bregma) were charted and collapsed onto a single sagittal plane. The location of each neuron was then defined by its x-y coordinates. To capture the aggregate spatial distribution, a geometric centroid of each neuron population was then determined to represent the center of mass in both x and y dimensions. Centroids were then used as the origin for the polar histograms. The size of each sector represents the relative neuron count as a function of direction. Histological and analysis procedures for projections from cortical neuron subtypes have been described previously (Hooks et al., 2018).

### Serial two-photon tomography

Serial two-photon tomography was used to map input to the GPe from the entire cortex. Imaging and analysis were performed as previously described (Kim et al., 2017). Two weeks after LVretro-Cre and CTb-488 injection, mouse brains were fixed as described above. Brains were then transferred to PBS and stored at 4 °C until imaged. Brains were embedded in 4% agarose in 0.05 M phosphate buffer and cross-linked in 0.2% sodium borohydrate solution (in PBS, pH 9.0–9.5). Each brain was imaged with a high-speed two-photon microscope with integrated vibratome (TissueVision) at 1 μm at both x-y resolution with 280 z-sections in every 50 μm. A 910-nm two-photon laser (Coherent) was used for CTb488 and tdTomato excitation. A dichroic mirror (Chroma) and band pass filters (Semrock) were used to separate green and red fluorescence signals. Emission signals were detected by GaAsP photomultiplier tubes (Hamamatsu). An automated, whole-brain cell counting and registration of the detected signal on a reference brain was applied as described before (Kim et al., 2017). The number of tdTomato^+^ neurons from each brain region was charted. The relative size of input to the GPe was calculated by normalizing the total number of tdTomato^+^ neurons in the entire brain of each sample.

### Visualized *ex vivo* electrophysiology

Mice in the age range postnatal day 55–90 were anesthetized with a ketamine-xylazine mixture and perfused transcardially with ice-cold aCSF containing the following (in mM): 125 NaCl, 2.5 KCl, 1.25 NaH_2_PO_4_, 2.0 CaCl_2_, 1.0 MgCl_2_, 25 NaHCO_3_, and 12.5 glucose, bubbled continuously with carbogen (95% O_2_ and 5% CO_2_). The brains were rapidly removed, glued to the stage of a vibrating microtome (Leica Instrument), and immersed in ice-cold aCSF. Parasagittal slices containing the dStr and the GPe were cut at a thickness of 240 μm and transferred to a holding chamber where they were submerged in aCSF at 37 °C for 30 min and returned to room temperature for recording. Slices were then transferred to a small-volume (∼0.5 ml) Delrin recording chamber that was mounted on a fixed-stage, upright microscope (Olympus). Neurons were visualized using differential interference contrast optics (Olympus), illuminated at 735 nm (Thorlabs), and imaged with a 60× water-immersion objective (Olympus) and a CCD camera (QImaging). Genetically-defined neurons were identified by somatic eGFP or tdTomato fluorescence examined under epifluorescence microscopy with a daylight (6,500 K) LED (Thorlabs) and appropriate filters (Semrock).

Recordings were made at room temperature (20–22 °C) with patch electrodes fabricated from capillary glass (Sutter Instrument) pulled on a Flaming-Brown puller (Sutter Instrument) and fire polished with a microforge (Narishige) immediately before use. Pipette resistance was typically ∼2–4 MΩ. For cell-attached and current-clamp recordings, the internal solution consisted of the following (in mM): 135 KMeSO_4_, 10 Na_2_phosphocreatine, 5 KCl, 5 EGTA, 5 HEPES, 2 Mg_2_ATP, 0.5 CaCl_2_, and 0.5 Na_3_GTP, with pH adjusted to 7.25–7.30 with KOH. The liquid junction potential for this internal solution was ∼7 mV and was not corrected. Stimulus generation and data acquisition were performed using an amplifier (Molecular Devices), a digitizer (Molecular Devices), and pClamp (Molecular Devices). For current-clamp recordings, the amplifier bridge circuit was adjusted to compensate for electrode resistance and was subsequently monitored. The signals were filtered at 1 kHz and digitized at 10 kHz. KMeSO_4_ and Na_2_-GTP were from ICN Biomedicals and Roche, respectively. All other reagents were obtained from Sigma-Aldrich.

For optogenetic experiments, blue excitation wavelength (peak, ∼450 nm) from two daylight (6,500 K) LEDs (Thorlabs) was delivered to the tissue slice from both a 60× water immersion objective and a 0.9 numerical aperture air condenser with the aid of 520 nm dichroic beamsplitters (Semrock). Light delivery was made at the site of electrophysiological recordings with a field of illumination of 500–700 µm in diameter. Paired-pulse optogenetic activation of terminals was at 20 Hz with a light duration of 2 ms.

### Experimental design and statistical analyses

General graphing and statistical analyses were performed with MATLAB (MathWorks), Prism (GraphPad), JASP (https://jasp-stats.org), and the R environment (https://www.r-project.org). Custom analysis codes are available on GitHub (https://github.com/chanlab). Sample size (*n* value) is defined by the number of observations (i.e., neurons, sections). When percentages are presented, *n* values represent only positive observations. No statistical method was used to predetermine sample size. All analyses are on the complete cases, not including imputation of any missing values. Data in the main text are presented as median values ± median absolute deviations (MADs) as measures of central tendency and statistical dispersion, respectively. Box plots are used for graphic representation of population data unless stated otherwise (Krzywinski and Altman, 2014; Streit and Gehlenborg, 2014). The central line represents the median, the box edges represent the interquartile ranges, and the whiskers represent 10–90^th^ percentiles. Normal distributions of data were not assumed. Individual data points were visualized for small sizes or to emphasize variability in the data. For each characteristic, a Kruskal–Wallis test was performed to detect an overall difference between neuron types. Lhx6^+^ neurons were considered together between all pairs of neuron types and evaluated using Dunn’s test, while differences between the Lhx6^+^_bright_ and Lhx6^+^_dim_ subtypes and all other cell types were evaluated using the Mann–Whitney *U* test. Unless < 0.0001, exact *P* values (two-tailed) are reported. Given that multiple tests were performed, in order to maintain an overall family-wise error rate of α = 0.05, the Bonferroni approach was used for each of the three sets of statistical tests. *K*-means analysis was performed in MATLAB with the number of clusters varying from two to six, considering spontaneous rate and CV_ISI_ as the clustering variables. Principal component analysis was performed in ClustVis (https://biit.cs.ut.ee/clustvis/) (Metsalu and Vilo, 2015); all cell-attached and whole-cell measurements were included in this analysis. Both rows and columns were clustered using correlation distance and average linkage. Electrophysiological measurements were centered and scaled. The median differences in spontaneous rate between different neuron types and the PV^+^ neurons and Npas1^+^ neurons and their 95% confidence intervals were estimated using the Estimation Stats application (https://www.estimationstats.com); five thousand bootstrap samples were taken; bias-corrected and accelerated (BC_a_) correction were applied to the resampling bootstrap distributions to adjust for both bias and skewness (Ho et al., 2019). Logistic regressions evaluating the correlation between the different variables and the Npas1^+^ neurons and PV^+^ neurons were also performed using the R programming language. Correlation matrix and network analysis were generated in JASP. Fruchterman-Reingold algorithm was used to iteratively compute the optimal placement of nodes.

## Results

### PV^+^ and Npas1^+^ neurons are distinct GPe neuron classes

Our laboratory and others have shown previously that the GPe contains two principal, near-exclusive classes of neurons distinguished by their respective expression of PV and Npas1 (Hegeman et al., 2016). However, the reported abundance of PV^+^ neurons across laboratories ranges from 30–60%, and their overlap with Npas1^+^ neurons within the GPe varies widely in the literature (Abdi et al., 2015; Abrahao and Lovinger, 2018; Dodson et al., 2015; Flandin et al., 2010; Mastro et al., 2014; Nobrega-Pereira et al., 2010; Oh et al., 2017; Saunders et al., 2015). As multiple mouse lines and well-characterized antibodies are available to study PV^+^ neurons, we sought to re-evaluate the abundance of PV^+^ GPe neurons and reconcile the inconsistencies. First, we examined PV immunoreactivity (PV-ir) across all GPe neurons using HuCD as a neuronal marker; we observed PV-ir in ∼50% of GPe neurons (49 ± 4%, *n =* 2,726 neurons, 19 sections) (Figure 1c). Note that the range observed within the PV population data (Figure 1c) mirrors the variance reported in the literature, suggesting that the discrepancy in the literature is likely of biological origin rather than methodological. Four PV antibodies were used throughout the study (see Table 2). All PV antibodies yielded similar results and were therefore pooled. While both PV-tdTom and PV-L-tdTom (PV-Cre;LSL-tdTomato) have been used consistently across laboratories for the identification of PV^+^ neurons, the utility of the PV-Flp line for studies of GPe neurons had not been established. To harness the potential of this mouse line, we crossed it with a Flp-reporter (FSF-tdTomato) line (see Materials and Methods). The resultant PV-F-tdTom (PV-Flp;FSF-tdTomato) line produced robust, cytoplasmic neuron labeling similar to that of the PV-L-tdTom and PV-tdTom lines. As expected, the PV-F-tdTom line showed prominent tdTomato expression (tdTomato^+^) not only in GPe PV^+^ neurons but also in PV^+^ neurons in the thalamic reticular nucleus (TRN) (Figure 1d and e) and cerebellum, as well as in PV^+^ interneurons throughout the cortex, hippocampus, and striatum.

**Figure 1.**
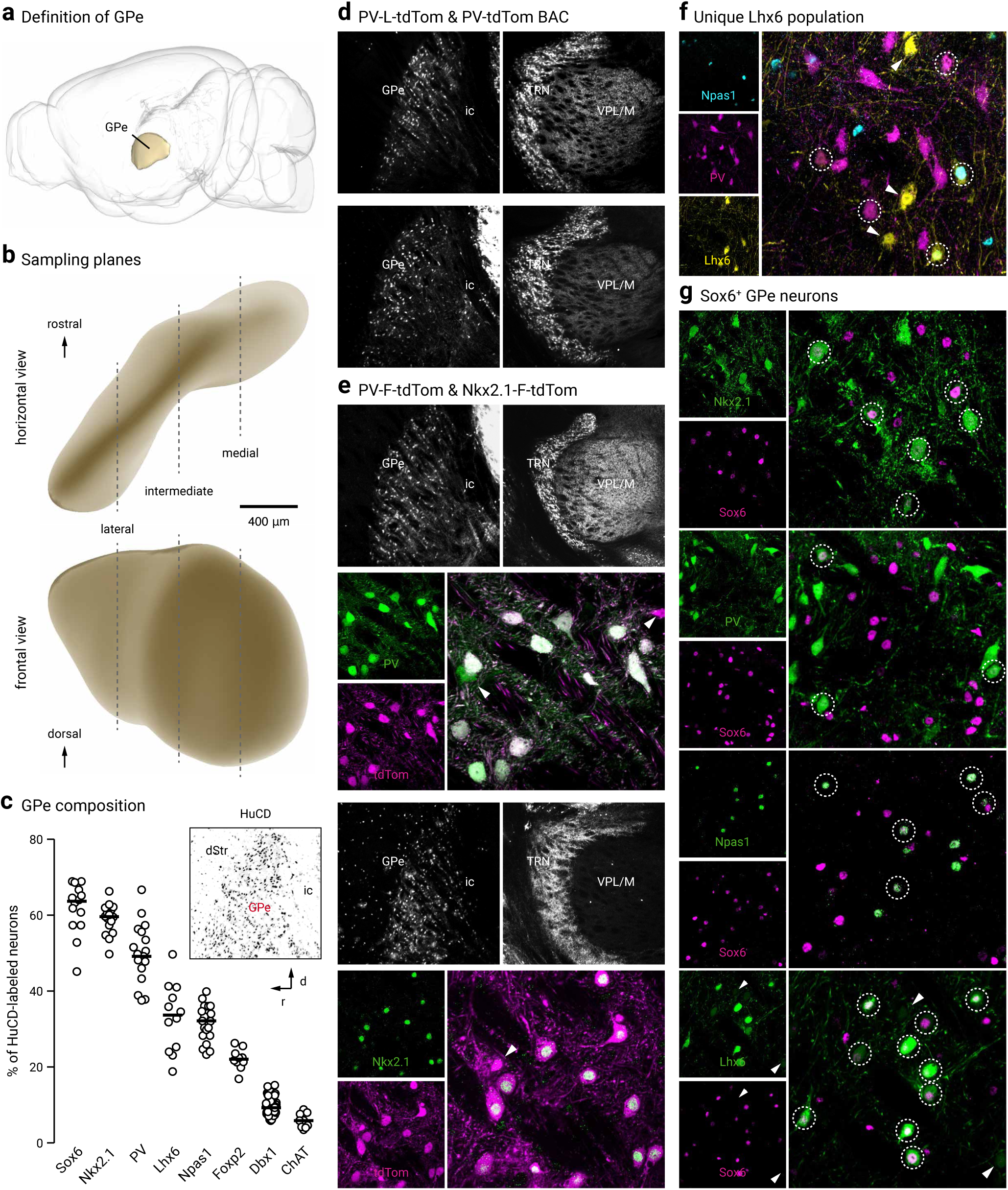
GPe neuron diversity. **a**. The location of the GPe in a mouse brain is illustrated (side view). **b**. GPe neurons at three different lateral, intermediate, and medial levels (∼2.5, 2.1, and 1.7 lateral from bregma) were sampled. **c**. Using HuCD as a neuronal marker, population data for the relative abundance of GPe neuron markers was determined. Each circle represents a section. Inset: low magnification confocal image of a sagittal brain section showing HuCD-labeling in the GPe with the dStr and ic defining the rostral and caudal borders, respectively. Note the low density of HuCD-labeled cells outside of the GPe. **d**. Low magnification confocal images of the GPe and TRN in PV-L-tdTom (PV-Cre;LSL-tdTomato, top) and PV-tdTom (bottom) mice. **e**. PV-F-tdTom (PV-Flp;FSF-tdTomato) and Nkx2.1-F-tdTom (Nkx2.1-Flp;FSF-tdTomato) were used in this study. The PV-F-tdTom (PV-Flp;FSF-tdTomato) transgenic mouse line produces faithful reporting of PV^+^ neurons and similar cytoplasmic neuron labeling as the PV-L-tdTom and PV-tdTom lines (as shown in **d**). Top, low magnification showing the PV-F-tdTom line produces prominent tdTomato expression (tdTomato^+^) in PV^+^ neurons in the TRN in addition to the GPe. To confirm the validity of the mouse line, tdTomato expression was compared against PV immunostaining. A higher magnification of the GPe shows nearly all tdTomato^+^ (magenta) neurons colocalize with PV-ir (green). Bottom, Nkx2.1-F-tdTom reliably captures neurons that arise from the Nkx2.1 lineage. Note that no cell bodies were found in the TRN (see also, Figure 5c). Double immunolabeling with tdTomato and Nkx2.1 demonstrated ∼90% colocalization. Arrowheads indicate neurons that do not colocalize. **f**. Triple immunostaining with PV, Npas1, and GFP on Lhx6-GFP brain sections confirms the existence of a Lhx6^+^-PV^−^-Npas1^−^ GPe population. Circles indicate Lhx6^+^ neurons that colocalize with either PV or Npas1. Arrowheads point to unique Lhx6^+^ neurons. Note that there are both bright and dim populations of Lhx6^+^ neurons. **g**. Sox6^+^ neurons express established GPe markers. Note that there are both bright and dim populations of Sox6^+^ neurons. Bottom, arrowheads indicate Lhx6^+^-Sox6^−^ neurons. Abbreviations: dStr = dorsal striatum; TRN = thalamic reticular nucleus; VPL/VPM = ventral posterior nucleus; ic = internal capsule.

Across the PV-tdTom, PV-L-tdTom, and PV-F-tdTom lines, the abundance of tdTomato^+^ neurons converged to ∼50% of the total GPe neuron population (PV-tdTom, 50 ± 9%, *n* = 571 neurons, 5 sections; PV-L-tdTom, 50 ± 13%, *n* = 848 neurons, 9 sections; PV-F-tdTom, 53 ± 15%, *n* = 859 neurons, 6 sections). Comparison of tdTomato expression with PV-ir demonstrated faithful reporting of PV^+^ neurons in all three transgenic lines (PV-tdTom, 100 ± 0%, *n* = 742 out of 747 neurons, 6 sections; PV-L-tdTom, 100 ± 0%, *n* = 1,380 out of 1,392 neurons, 12 sections; PV-F-tdTom, 100 ± 0%, *n* = 930 out of 1,023 neurons, 6 sections).

The Npas1-Cre-tdTom line was previously generated in our laboratory, and it labels roughly 90% (88 ± 7%, *n* = 505 neurons) of neurons that endogenously express Npas1 in the GPe (Hernandez et al., 2015). As this mouse line does not fully capture all Npas1^+^ neurons, we focused our quantifications on Npas1-immunoreactivity (Npas1-ir) with our previously characterized Npas1 antibodies (Hernandez et al., 2015). We found that Npas1^+^ neurons make up ∼30% (32 ± 4%, *n =* 3,361 neurons, 21 sections) of all GPe neurons. This result is consistent with our previous quantification of 27% of the entirety of the GPe. A re-evaluation of the overlap between PV and Npas1 confirmed our prior observation (Hernandez et al., 2015) that these two markers are expressed in almost completely segregated neuron populations. We observed a very low level of overlap across lateromedial GPe levels (2 ± 2%, *n* = 96 out of 1,777 neurons, 12 sections), with the most overlap observed at the intermediate level (lateral: 1 ± 1%, *n* = 17 neurons, 3 sections; intermediate: 3 ± 1%, *n* = 68 neurons, 5 sections; medial: 0 ± 0%, *n* = 11 neurons, 4 sections). This is slightly higher than our previous estimate and may be related to our previous reliance on the Npas1 mouse line for quantification (Hernandez et al., 2015). Our data are at odds with Abrahao and Lovinger, 2018, which shows a much higher (up to 12.6%) overlap between PV and Npas1 expression in the GPe. This discrepancy can be attributed to differences in immunolabeling protocols, quantification methods, and the age of mice used in the experiments.

As determined by immunolabeling, a small population of neurons expressed a cholinergic neuron marker, choline acetyltransferase (ChAT) (6 ± 2%, *n* = 267 neurons, 9 sections), a finding that is consistent with our previous reports (Gritti et al., 2006; Hernandez et al., 2015; Nobrega-Pereira et al., 2010) (see Table 3). This leaves around 15% of the GPe population that is unidentified by PV, Npas1, or ChAT.

**Table 3.** Quantification of GPe neurons

|  | Molecularly-defined neuron subtypes |  |  |  |  |  |  |  |  |  |  |  |  |  |  |  |
| --- | --- | --- | --- | --- | --- | --- | --- | --- | --- | --- | --- | --- | --- | --- | --- | --- |
|  | Sox6<sup>+</sup> |  | Nkx2.1<sup>+</sup> |  | PV<sup>+</sup> |  | <sup>a</sup>Lhx6<sup>+</sup> |  | Npas1<sup>+</sup> |  | Foxp2<sup>+</sup> |  | Dbx1<sup>+</sup> |  | ChAT<sup>+</sup> |  |
|  | Median<br>±<br>MAD | *n*<br>(neurons;<br>sections) | Median<br>±<br>MAD | *n*<br>(neurons;<br>sections) | Median<br>±<br>MAD | *n*<br>(neurons;<br>sections) | Median<br>±<br>MAD | *n*<br>(neurons;<br>sections) | Median<br>±<br>MAD | *n*<br>(neurons;<br>sections) | Median<br>±<br>MAD | *n*<br>(neurons;<br>sections) | Median<br>±<br>MAD | *n*<br>(neurons;<br>sections) | Median<br>±<br>MAD | *n*<br>(neurons;<br>sections) |
| Total GPe neurons (%) | 63.7 ± 4.0 | 4,681; 14 | 59.6 ± 2.3 | 5,342; 16 | 49.2 ± 4.3 | 2,726; 19 | 33.7 ± 7.5 | 2,533; 12 | 32.1 ± 3.9 | 3,361; 21 | 22.1 ± 1.1 | 1,273; 10 | 9.3 ± 1.3 | 3,002; 61 | 5.9 ± 1.7 | 267; 9 |
| Coexpression (%) |  |  |  |  |  |  |  |  |  |  |  |  |  |  |  |  |
| Sox6 | – | – | 68.2 ± 0.5 | 681; 3 | 52.5 ± 8.4 | 1,675; 15 | <sup>d</sup>75.5 ± 7.2 | 2,346; 15 | 93.4 ± 5.6 | 1,999; 14 | 67.0 ± 4.3 | 206; 3 | 51.6 ± 8.2 | 442; 15 | 0.0 ± 0.0 | 0; 3 |
| Nkx2.1 | 58.2 ± 0.3 | 681; 3 | – | – | 83.2 ± 3.5 | 1,105; 7 | 78.4 ± 8.0 | 1,469; 9 | 32.0 ± 1.4 | 383; 6 | 0.0 ± 0.0 | 1; 3 | 77.8 ± 10.2 | 329; 9 | – | – |
| PV | 36.4 ± 3.2 | 1,675; 15 | 70.8 ± 3.0 | 1,105; 7 | – | – | <sup>b</sup>28.1 ± 7.3 | 654; 12 | 2.7 ± 2.7 | 96; 12 | – | – | 72.0 ± 8.3 | 493; 23 | – | – |
| Lhx6 | 47.0 ± 6.7 | 2,346; 15 | 40.6 ± 7.4 | 1,469; 9 | 24.1 ± 9.6 | 654; 12 | – | – | 40.7 ± 4.6 | 818; 12 | 1.3 ± 1.3 | 8; 3 | – | – | 50.0 ± 3.6 | 48; 3 |
| Npas1 | 45.4 ± 5.7 | 1,999; 14 | 17.3 ± 2.3 | 383; 6 | 2.0 ± 2.0 | 96; 12 | <sup>c</sup>35.2 ± 5.1 | 818; 12 | – | – | 79.3 ± 3.8 | 566; 6 | 9.5 ± 5.2 | 72; 14 | – | – |
| Foxp2 | 34.1 ± 5.7 | 206; 13 | 0.0 ± 0.0 | 1; 3 | – | – | 1.9 ± 0.5 | 8; 3 | 56.8 ± 4.2 | 566; 6 | – | – | 3.6 ± 1.9 | 13; 10 | – | – |
| Dbx1 | 10.4 ± 3.2 | 411; 15 | 12.0 ± 1.3 | 329; 9 | 13.6 ± 3.0 | 493; 23 | – | – | 3.3 ± 1.8 | 72; 14 | 1.5 ± 0.6 | 13; 10 | – | – | 10.6 ± 4.2 | 34; 9 |
| ChAT | 0.0 ± 0.0 | 0; 3 | – | – | – | – | <sup>g</sup>6.7 ± 0.6 | 48; 3 | – | – | – | – | 7.4 ± 3.3 | 34; 9 | – | – |

<sup>a</sup>Lhx6<sup>+</sup> neuron composition: 28% <sup>b</sup>(PV<sup>+</sup>) + 35% <sup>c</sup>(Npas1<sup>+</sup>) + 37% <sup>e+f</sup>(Lhx6<sup>+</sup>-PV<sup>-</sup>-Npas1<sup>-</sup>) = 100%
<sup>e</sup>Lhx6<sup>+</sup>-PV<sup>-</sup>-Npas1<sup>-</sup>-Sox6<sup>-</sup>: 100% – d = 24%
<sup>f</sup>Lhx6<sup>+</sup>-PV<sup>-</sup>-Npas1<sup>-</sup>-Sox6<sup>+</sup>: d – b – c = 13%
<sup>g</sup>ChAT<sup>+</sup> neurons can be a portion of Lhx6<sup>+</sup>-PV<sup>-</sup>-Npas1<sup>-</sup>-Sox6<sup>-</sup> neurons
**Lhx6<sup>+</sup>-PV<sup>+</sup> and Lhx6<sup>+</sup>-Npas1<sup>+</sup> neurons also express Sox6**
Abbreviation: MAD = median absolute deviation.

### Nkx2.1-F-tdTom mice label prototypic GPe neurons

Based on our previous data showing that Lhx6^+^ neurons overlap substantially with both PV^+^ neurons and Npas1^+^ neurons (Hernandez et al., 2015), we hypothesized that a subset of Lhx6^+^ neurons may correspond to the 15% of GPe neurons that are PV^−^ and Npas1^−^ (and ChAT^−^) (see Figure 1f). However, the use of the Lhx6-GFP mouse line has resulted in highly inconsistent results across laboratories (Hegeman et al., 2016).

To glean insights into an improved GPe neuron classification, we systematically examined the GPe expression of Nkx2.1 and Sox6, signaling molecules that are upstream and downstream of Lhx6, respectively. This was motivated by the observation that Nkx2.1, Lhx6, and Sox6 work in concert to control the fate of forebrain GABAergic neurons (Batista-Brito et al., 2009; Du et al., 2008; Jaglin et al., 2012). Consistent with the fact that Nkx2.1 plays a crucial role in GPe development (Flandin et al., 2010; Nobrega-Pereira et al., 2010; Rubenstein et al., 1998; Sussel et al., 1999), Nkx2.1 immunolabeling revealed that ∼60% (60 ± 2%, *n* = 5,342 neurons, 16 sections) of GPe neurons are Nkx2.1-expressing (Nkx2.1^+^) (Figure 1c and e, Table 3). In keeping with our previous analysis of Lhx6 and Foxp2 (Hernandez et al., 2015), we observed no overlap between Nkx2.1^+^ neurons and Foxp2-expressing (Foxp2^+^) neurons in wild-type brain sections (*n* = 1 out of 1,131 Nkx2.1^+^ neurons, 3 sections; *n* = 1 out of 414 Foxp2^+^ neurons, 3 sections). Subsequent immunohistological analysis of Nkx2.1 antibody labeling with established GPe markers revealed that the majority of Nkx2.1^+^ neurons are PV^+^ (71 ± 3%, *n =* 1,105 neurons, 7 sections), while a smaller subset are Npas1^+^ (17 ± 2%, *n =* 383 neurons, 6 sections) or Lhx6^+^ (41 ± 7%, *n =* 1,469 neurons, 9 sections) (Table 3). Importantly, Nkx2.1^+^ neurons are only ∼80% (83 ± 4%, *n* = 1,105 neurons, 7 sections) of PV^+^ neurons, consistent with previous observations (Xu et al., 2008). Furthermore, Nkx2.1^+^ neurons represent ∼80% (78 ± 8%, *n* = 1,469 neurons, 9 sections) of Lhx6^+^ neurons and only a subset of Npas1^+^ neurons (32 ± 1%, *n* = 383 neurons, 6 sections). More importantly, with triple immunostaining, we found that nearly all Npas1^+^-Nkx2.1^+^ neurons are Lhx6^+^ (90 ± 5%, *n* = 184 out of 204 neurons, 3 sections). Accordingly, the Npas1^+^-Nkx2.1^+^ and Npas1^+^-Lhx6^+^ populations of Npas1^+^ neurons are near-identical (33 ± 5%, *n* = 204 out of 609 neurons, 3 sections and 32 ± 3%, *n* = 233 out of 609 neurons, 3 sections, respectively) (data not shown). This finding is in line with what we have previously described (Hernandez et al., 2015).

To effectively identify all neurons derived from the Nkx2.1 lineage, we made a Nkx2.1-F-tdTom (Nkx2.1-Flp;FSF-tdTomato) genetic cross, which yielded robust tdTomato labeling (tdTomato^+^) in the GPe. In addition to tdTomato^+^ neurons, we observed tdTomato^+^ glia in the GPe that were distinguished from neurons based on their morphology. To assess the cumulative recombination events in Nkx2.1-F-tdTom, we compared tdTomato expression to Nkx2.1 immunolabeling. Immunoreactivity on Nkx2.1-F-tdTom brain sections with tdTomato, Nkx2.1, and HuCD antibodies revealed ∼90% (89 ± 11%, *n* = 795 neurons, 3 sections) of tdTomato^+^ neurons colocalized with Nkx2.1^+^ neurons (Figure 1e). While we observed significant tdTomato^+^ labeling throughout the entire cortex, Nkx2.1 immunoreactivity was absent. These data corroborate previous findings that Nkx2.1 is downregulated in cortical interneurons in adult mice (Butt et al., 2008; Nobrega-Pereira et al., 2008) and that the majority of GPe neurons that are derived from the Nkx2.1 lineage maintain Nkx2.1 expression into adulthood. Similarly, we observed no tdTomato^+^ or Nkx2.1^+^ cell bodies in regions caudal to the forebrain, such as the TRN (Figure 1e), STN, and SNr (data not shown).

### Sox6 delineates GPe neuron subtypes

Previous literature demonstrates that the transcription factor Sox6 is present in most, if not all, medial ganglionic eminence (MGE)-derived neurons in the mature brain (Azim et al., 2009; Batista-Brito et al., 2009). Moreover, Sox6 and Lhx6 colocalize extensively within the MGE (Batista-Brito et al., 2009), consistent with the function of Sox6 as a downstream signaling partner of Lhx6 (Azim et al., 2009; Batista-Brito et al., 2009). In view of these findings, we set out to examine the Sox6 expression pattern in the GPe (Figure 1g). Sox6 immunolabeling revealed ∼65% (64 ± 4%, *n =* 4,681 neurons, 14 sections) of GPe neurons are Sox6-expressing (Sox6^+^). We first investigated Sox6 expression in Nkx2.1^+^ neurons; due to incompatibility of the Nkx2.1 and Sox6 antibodies, we relied on Sox6 immunolabeling in Nkx2.1-F-tdTom brain sections. We found substantial overlap between tdTomato^+^ neurons and Sox6^+^ neurons (68 ± 1% out of Nkx2.1, *n* = 681 neurons, 3 sections; 58 ± 0% out of Sox6, *n* = 681 neurons, 3 sections) (Table 3). Although both Sox6 and Nkx2.1 are expressed in ∼60–65% of GPe neurons, they do not represent the same pool of neurons, as their expression overlaps differently with other GPe neuron markers (Table 3). Nearly all Npas1^+^ neurons are Sox6^+^ (93 ± 6%, *n =* 1,999 neurons, 14 sections), while only half of the PV^+^ population expresses Sox6 (53 ± 8%, *n =* 1,675 neurons, 15 sections). As mentioned above, Nkx2.1^+^ neurons account for ∼32% of Npas1^+^ neurons and ∼83% of PV^+^ neurons.

Next, we examined the relationship between the Sox6 and Lhx6 GPe neuron populations, as this may yield important insights into targeting the otherwise difficult-to-identify Lhx6^+^-PV^−^-Npas1^−^ population. As we were unsuccessful in labeling endogenous Lhx6 protein with multiple Lhx6 antibodies (Santa Cruz SC-98607, Santa Cruz SC-271433, NeuroMab 73-241, and a custom antibody from Dr. John Rubenstein), we used the GFP expression in Lhx6-GFP mice as a proxy for Lhx6 expression. Overall, Lhx6^+^ neurons represent a third of the GPe (34 ± 8%, *n* = 2,533 neurons, 12 sections); approximately 30% of these neurons express PV (28 ± 7%, *n* = 654 neurons, 12 sections), 35% express Npas1 (35 ± 5%, *n* = 818 neurons, 12 sections), and 7% express ChAT (7 ± 1%, *n* = 48 neurons, 3 sections). These percentages confirm our previous reports (Hernandez et al., 2015). To determine the relationship between Sox6 expression within the Lhx6^+^, PV^+^, and Npas1^+^ populations, immunolabeling on Lhx6-GFP sections showed that Sox6 was expressed in ∼75% (76 ± 7%, *n* = 2,346 neurons, 15 sections) of Lhx6^+^ neurons (Figure 1g) as well as in all PV^+^ neurons and Npas1^+^ neurons that were Lhx6^+^ (Figure 2a). Note that similar to Lhx6, there were bright and dim populations of Sox6. We did not observe a relationship between Sox6 fluorescence levels and expression of PV and Npas1. Consistent with our previous observations (Hernandez et al., 2015), we observed negligible overlap between Lhx6^+^ and Foxp2^+^ GPe neurons (8 out of 1,882 neurons, 3 sections).

**Figure 2.**
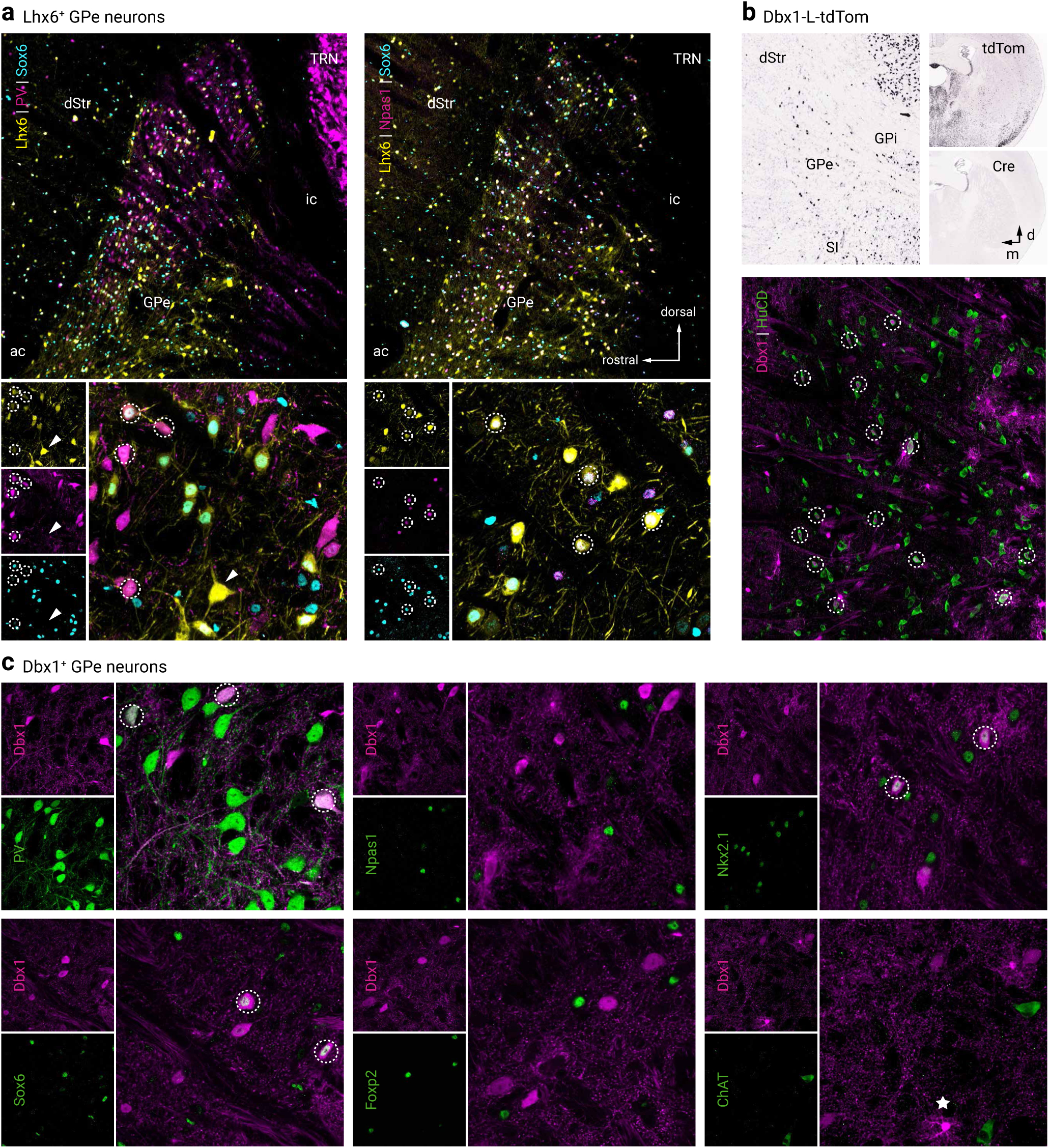
Lhx6^+^ and Dbx1^+^ GPe neurons colocalize with established GPe markers. **a.** Left, low and high magnification images of PV^+^, Sox6^+^, and Lhx6^+^ (GFP^+^) GPe neurons. Right, low and high magnification images of Npas1^+^, Sox6^+^, and Lhx6^+^ (GFP^+^) GPe neurons. **b**. *In situ* hybridization signals from Dbx1-L-tdTom mouse line for tdTomato^+^ and Cre^+^ in the adult GPe and neighboring areas (left). Note the widespread tdTomato^+^ across brain areas (top right) resulted from the cumulative recombination from early developmental stages in spite of the absence of Cre^+^ expression in adult (bottom right). Data are adopted from Allen Brain Atlas. The Dbx1-L-tdTom mouse line labeled Dbx1^+^ (tdTomato^+^) neurons (HuCD, magenta) and glia in the GPe. **c**. Dbx1^+^ GPe neurons colocalized with established GPe markers and are largely PV^+^. Note that there was no overlap between Dbx1 and Foxp2. Circles indicate colocalization, star (bottom right) presents an example of astrocytic labeling in the Dbx1-L-tdTom mouse line. Abbreviations: dStr = dorsal striatum; GPi = internal globus pallidus; SI = substantia innominata; TRN = thalamic reticular nucleus; ac = anterior commissure; ic = internal capsule.

Taken together, our data describe a Lhx6^+^ population (∼37%) that is devoid of both PV and Npas1 expression and accounts for ∼13% of the entire GPe. A portion of this Lhx6^+^-PV^−^-Npas1^−^ population does not express Sox6 and represents roughly a quarter of Lhx6^+^ neurons (24 ± 6%, *n* = 938 neurons, 15 sections) or 8% of all GPe neurons, whereas the Lhx6^+^-PV^−^-Npas1^−^ population that does express Sox6 accounts for only ∼4% of the entire GPe. Given that 7% of Lhx6^+^ neurons express ChAT, and we observed no overlap between ChAT and Sox6 (0 out of 1,674 neurons, 3 sections), we were able to pinpoint the molecular identity of the entirety of Lhx6^+^ neurons (see footnote in Table 3). Importantly, the Lhx6^+^-Nkx2.1^+^ population is likely equivalent to the Lhx6^+^-Sox6^+^ population, as both Nkx2.1 and Sox6 overlap similarly with Lhx6^+^ (Nkx2.1 out of Lhx6, 78 ± 8%, *n* = 1,469 neurons, 9 sections; Sox6 out of Lhx6, 76 ± 7%, *n* = 2,346 neurons, 15 sections), and the Lhx6^+^-Nkx2.1^+^ and Lhx6^+^-Sox6^+^ populations are similarly abundant within the entire GPe (28 ± 7% and 32 ± 6%, respectively). However, with our present tools, we cannot confirm the assertion that these two subtypes are identical.

### Dbx1-L-tdTom mice label a heterogenous GPe neuron subset

Since Lhx6^+^-Sox6^−^ neurons constitute a substantial fraction (8%) of the GPe, we sought to identify an additional expression marker for this population. Neurons from the Dbx1 lineage originate from the preoptic area (PoA) and are known to populate the GPe (Nobrega-Pereira et al., 2010). Considering Lhx6 is expressed in postmitotic neurons derived from both the MGE and the PoA (Du et al., 2008; Fogarty et al., 2007), we set out to investigate if neurons from the Dbx1 lineage correspond to Lhx6^+^-Sox6^−^ GPe neurons. Accordingly, we identified neurons that arise from the Dbx1 lineage and determined their relationship with the Lhx6^+^-Sox6^−^ GPe subset by using a Dbx1-L-tdTom (Dbx1-Cre;LSL-tdTomato) cross, which produced robust tdTomato expression in the GPe. In addition to neuronal labeling, tdTomato^+^ glia were present within the GPe and densely populated the ventral GPe border. For simplicity, we refer to tdTomato^+^ neurons in Dbx1-L-tdTom mice as Dbx1^+^ neurons. Using HuCD as a neuronal marker (Figure 2b), we determined that Dbx1^+^ neurons account for ∼9% of the entire GPe population (9 ± 1%, *n* = 3,002 neurons, 61 sections). Immunohistological analysis of Dbx1^+^ neurons showed substantial co-expression with Nkx2.1 (78 ± 10%, *n =* 329 neurons, 9 sections) and Sox6 (52 ± 8%, *n =* 442 neurons, 15 sections). Furthermore, Dbx1^+^ neurons are primarily PV^+^ (72 ± 8%, *n* = 493 neurons, 23 sections) and to a lesser extent Npas1^+^ (10 ± 5%, *n =* 72 neurons, 14 sections), ChAT^+^ (7 ± 3%, *n =* 34 neurons, 9 sections), and Foxp2^+^ (4 ± 2%, *n =* 13 neurons, 10 sections) (Figure 2c). To summarize, we found Dbx1^+^ neurons do not correspond to the Lhx6^+^-Sox6^−^ unique GPe subset. While both populations were of similar abundance in the GPe, they varied in their co-expression of PV, Npas1, and Sox6. Expression of Lhx6 in Dbx1^+^ neurons was not examined in this study for technical reasons; however, due to the close functional relationship between Nkx2.1, Sox6, and Lhx6, it can be inferred that Lhx6 represents a substantial fraction of Dbx1^+^ neurons.

### Dbx1^+^-PV^+^ neurons exhibit canonical PV^+^ neuron projection patterns

Although Dbx1^+^ neurons account for only ∼10% of GPe neurons, it is possible that they target a unique area and serve an important function. As Cre is not expressed in adult Dbx1-Cre mice (Bielle et al., 2005) (Figure 2b), we do not have true genetic access to this neuron population. Rather than relying on standard Cre-inducible viral approaches, Alexa-conjugated cholera toxin b (CTb), a widely-used retrograde tracer, was used to map the axonal projection patterns of Dbx1^+^ neurons. Given the majority of Dbx1^+^ neurons are PV^+^, we first targeted the STN, the principal recipient of PV^+^ GPe input, in our connectome survey (Hegeman et al., 2016; Hernandez et al., 2015). As expected, we found Dbx1^+^ neurons project primarily to the STN (23 ± 9%, *n* = 94 neurons, 9 sections) (Figure 3a and b, Table 4). Additionally, but to a much lesser extent, we found Dbx1^+^ neurons project to the substantia nigra (SN, 2 ± 0%, *n =* 20 neurons, 9 sections). These numbers are likely an underestimation due to incomplete coverage of the target areas (i.e., STN and SN) with these injections. Importantly, although ∼10% of Dbx1^+^ GPe neurons co-express Npas1, we observed no projections to the dorsal striatum (0 out of 42 CTb^+^ neurons, 17 sections) (Table 4). The negative results were not due to poor labeling efficiency as neighboring Dbx1^−^ neurons were evidently labeled in the same section. To examine if Dbx1^+^ neurons account for the recently described non-cholinergic, cortically-projecting GPe neurons (Ahrlund-Richter et al., 2019; Chen et al., 2015; Saunders et al., 2015; Schwarz et al., 2015; Sun et al., 2019; Van der Kooy and Kolb, 1985), CTb was injected into various cortical regions of Dbx1-L-tdTom mice. Systematic analysis of multiple cortical areas including the somatomotor (MO, *n* = 4 mice), somatosensory (SS, *n* = 2 mice), anterior cingulate (AC, *n* = 1 mouse), agranular (AG) and orbital (ORB, *n* = 2 mice) cortices revealed Dbx1^+^ neurons do not project to any of these cortical regions (0 out of 52 CTb^+^ neurons, 9 sections). The presence of CTb^+^ neurons within the GPe confirmed that the negative results were not due to failed injections (see below).

**Figure 3.**
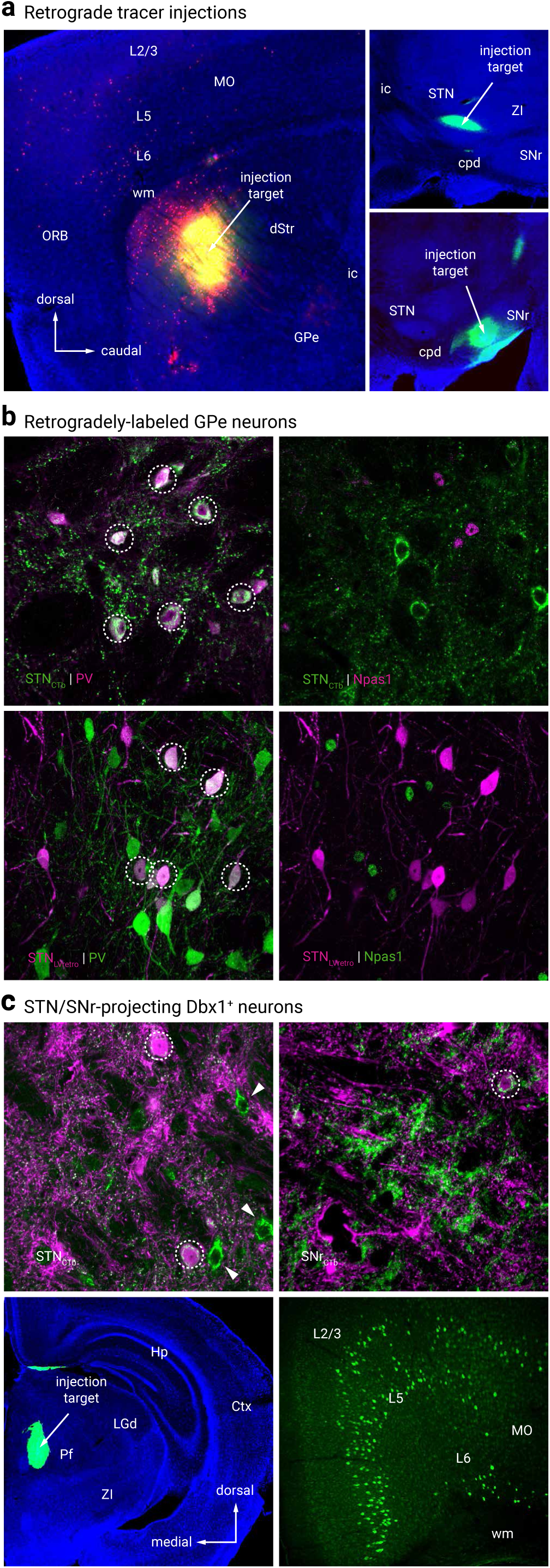
Retrograde tracing analysis. **a.** Representative injection sites from retrograde tracing connectome analysis. CTb (green) with or without LVretro-Cre (LV, red) was injected into dStr (top left), STN (top right), SNr (bottom right) and mounted with DAPI (blue) to visualize cytoarchitecture. **b**. Retrograde labeling of GPe-STN neurons with both CTb and LV tracing techniques. CTb (top left, green) labeled and LV (bottom left, magenta) GPe neurons from STN injection are primarily PV^+^ and did not colocalize with Npas1 immunostaining (top and bottom right). **c**. Retrograde labeling in Dbx1-L-tdTom mice shows Dbx1^+^ neurons (magenta) project to STN (top left) and SNr (top right) as indicated by colocalization with CTb (green). Circles denote colocalization. Arrowheads denote CTb^+^ STN projecting neurons that lack expression of Dbx1. Bottom, coronal view of a representative injection to the PF (left) along with expected positive cortical fluorescence (MO, right). No fluorescence was observed in the GPe. Abbreviations: dStr = dorsal striatum; Ctx = cortex; GPe = external globus pallidus; GPi = internal globus pallidus; Hp = hippocampus; Pf = parafascicular nucleus; LGd = lateral geniculate, dorsal; MO = somatomotor cortex; ORB = orbital cortex; SI = substantia innominata; SNr = substantia nigra pars reticulata; STN = subthalamic nucleus; TRN = thalamic reticular nucleus; ZI = zona incerta; ac = anterior commissure; cpd = cerebral peduncle; ic = internal capsule; wm = white matter.

**Table 4.** Quantification of retrogradely-labeled GPe neurons

| LVretro-cre | Injection targets |  |  |  |  |  |  |  |  |  |  |  |
| --- | --- | --- | --- | --- | --- | --- | --- | --- | --- | --- | --- | --- |
|  | STN |  |  | SN |  |  | dStr |  |  | Ctx (ORB, MO, SS) |  |  |
|  | Median<br>±<br>MAD | *n* (neurons) | *n* (sections) | Median<br>±<br>MAD | *n* (neurons) | *n* (sections) | Median<br>±<br>MAD | *n* (neurons) | *n* (sections) | Median<br>±<br>MAD | *n* (neurons) | *n* (sections) |
| Total tdTomato<sup>+</sup> | – | 750 | 18 | – | 127 | 12 | – | 406 | 15 | – | 293 | 24 |
| PV<sup>+</sup> | 75.6 ± 6.7 | 278 | 9 | 66.7 ± 26.2 | 51 | 9 | 11.5 ± 4.8 | 15 | 6 | 0.0 ± 0.0 | 3 | 8 |
| PV<sup>−</sup> | 20.0 ± 8.7 | 74 | 9 | 28.6 ± 25.3 | 18 | 9 | 88.5 ± 4.8 | 114 | 6 | 100.0 ± 0.0 | 62 | 11 |
| Npas1<sup>+</sup> | 15.6 ± 6.5 | 45 | 9 | 15.8 ± 6.4 | 7 | 3 | 75.0 ± 8.3 | 200 | 9 | 25.0 ± 6.3 | 41 | 13 |
| Npas1<sup>−</sup> | 79.1 ± 16.1 | 247 | 9 | 84.2 ± 6.4 | 51 | 3 | 25.0 ± 8.3 | 77 | 9 | 68.8 ± 11.3 | 106 | 13 |
| Npas1<sup>+</sup>-Nkx2.1<sup>+</sup> | – | – | – | – | – | – | – | – | – | 20.0 ± 6.7 | 8 | 3 |
| Nkx2.1<sup>+</sup> | – | – | – | – | – | – | – | – | – | 32.3 ± 5.8 | 20 | 6 |
| ChAT<sup>+</sup> | – | – | – | – | – | – | – | – | – | 50.0 ± 0.0 | 19 | 3 |
| ChAT<sup>−</sup> | – | – | – | – | – | – | – | – | – | 50.0 ± 0.0 | 23 | 3 |
| Foxp2<sup>+</sup> | – | – | – | – | – | – | – | – | – | 0.0 ± 0.0 | 0 | 3 |
| CTb | STN |  |  | SN |  |  | dStr |  |  | Ctx (ORB, MO, SS) |  |  |
|  | Median<br>±<br>MAD | *n* (neurons) | *n* (sections) | Median<br>±<br>MAD | *n* (neurons) | *n* (sections) | Median<br>±<br>MAD | *n* (neurons) | *n* (sections) | Median<br>±<br>MAD | *n* (neurons) | *n* (sections) |
| Total CTb<sup>+</sup> | – | 592 | 12 | – | 173 | 15 | – | 42 | 17 | – | 116 | 21 |
| PV<sup>+</sup> | 72.6 ± 11.1 | 263 | 9 | 33.3 ± 11.1 | 14 | 9 | 0.0 ± 0.0 | 1 | 8 | 0.0 ± 0.0 | 0 | 3 |
| PV<sup>−</sup> | 27.5 ± 10.6 | 99 | 9 | 66.7 ± 11.1 | 52 | 9 | 100.0 ± 0.0 | 18 | 8 | 100.0 ± 0.0 | 6 | 3 |
| Npas1<sup>+</sup> | 6.4 ± 4.4 | 10 | 3 | 18.6 ± 8.3 | 19 | 6 | 100.0 ± 0.0 | 18 | 9 | 40.4 ± 9.7 | 19 | 6 |
| Npas1<sup>−</sup> | 93.7 ± 4.4 | 130 | 3 | 81.4 ± 8.3 | 68 | 6 | 0.0 ± 0.0 | 5 | 9 | 60.0 ± 10.0 | 27 | 6 |
| Dbx1<sup>+</sup> | 22.6 ± 9.4 | 94 | 9 | 2.2 ± 0.3 | 20 | 9 | 0.0 ± 0.0 | 0 | 17 | 0.0 ± 0.0 | 0 | 9 |
| PV<sup>+</sup>-Dbx1<sup>+</sup> | 11.8 ± 11.8 | 56 | 9 | 5.6 ± 5.6 | 10 | 15 | 0.0 ± 0.0 | 0 | 8 | 0.0 ± 0.0 | 0 | 3 |
| PV<sup>−</sup>-Dbx1<sup>+</sup> | 0.0 ± 0.0 | 7 | 9 | 0.0 ± 0.0 | 6 | 15 | 0.0 ± 0.0 | 0 | 8 | 0.0 ± 0.0 | 0 | 3 |
| Npas1<sup>+</sup>-Dbx1<sup>+</sup> | 0.0 ± 0.0 | 0 | 3 | 0.0 ± 0.0 | 2 | 9 | 0.0 ± 0.0 | 0 | 9 | 0.0 ± 0.0 | 0 | 6 |
| Npas1<sup>−</sup>-Dbx1<sup>+</sup> | 14.6 ± 2.1 | 27 | 3 | 0.0 ± 0.0 | 6 | 9 | 0.0 ± 0.0 | 0 | 9 | 0.0 ± 0.0 | 0 | 6 |

Abbreviations: Ctx = cortex, dStr = dorsal striatum, MAD = median absolute deviation, MO = somatomotor area, ORB = orbital area, SN = substantia nigra, SS = somatosensory area, STN = subthalamic nucleus.

To confirm that the findings from retrograde tracing were not just due to obscure biases with the CTb, a pseudotyped-lentivirus was used for retrograde delivery of Cre recombinase (LVretro-Cre) (Knowland et al., 2017) in LSL-tdTomato mice. Unlike CTb, this strategy gives more robust, unambiguous cytoplasmic tdTomato expression. CTb and LVretro-Cre were injected into various known areas that receive GPe input, including dStr, STN, SNr, parafascicular nucleus (PF) (Kita, 2007). We observed comparable labeling patterns with LVretro-Cre and CTb injections into the dStr (Npas1^+^_LVretro_: 75 ± 8%, *n* = 200 neurons, 9 sections; Npas1^+^_CTb_: 100 ± 0%, *n* = 18 neurons, 9 sections) and STN (PV^+^_LVretro_: 76 ± 7%, *n* = 278 neurons, 9 sections; PV^+^_CTb_: 73 ± 11%, *n* = 263 neurons, 9 sections). A different scenario was observed with SNr injections—there was not a predominant neuron class that projected to the SNr (PV^+^_LVretro_: 67 ± 26%, *n* = 51 neurons, 9 sections; PV^+^_CTb_: 33 ± 11%, *n* = 14 neurons, 9 sections) (Figure 3c, Table 4). Such a discrepancy between LVretro-Cre and CTb observed in the SNr data could be attributed to a number of factors including complex topographical organization of the GPe projection and CTb spreading into areas that are adjacent to the SNr and receive Npas1^+^ input (e.g., substantia nigra pars compacta). Furthermore, LVretro-Cre and CTb injections into the PF yielded labeling throughout the cortex, most prominently in the MO and AG regions (Figure 3c), consistent with prior observations (Mandelbaum et al., 2019; Sherman, 2016). However, contrary to reports of a GPe-PF connection from other laboratories (Mastro et al., 2014), we observed no labeling in the GPe with these injections (LVretro-Cre, 4 mice; CTb, 9 mice). It is possible that CTb and LVretro-Cre both have low labeling efficiencies, making it difficult to conclude the absence of sparse projections.

### Cortex and GPe are reciprocally connected

While we found no Dbx1^+^ neurons projecting to the cortex, we identified a subset of GPe neurons that are cortex-projecting. CTb-based tracing revealed primary somatosensory (SSp, aka S1), primary somatomotor (MOp, aka M1), AG and ORB cortices as the primary targets of cortical-projecting GPe neurons (CTb, *n* = 116 neurons, 21 sections). To confirm these regions as the primary targets, we injected LVretro-Cre into frontal cortical regions of LSL-tdTomato mice. As expected, we observed a population of retrogradely labeled GPe neurons, i.e. tdTomato^+^ (LV, *n* = 293 neurons, 24 sections) (Figure 4b).

**Figure 4.**
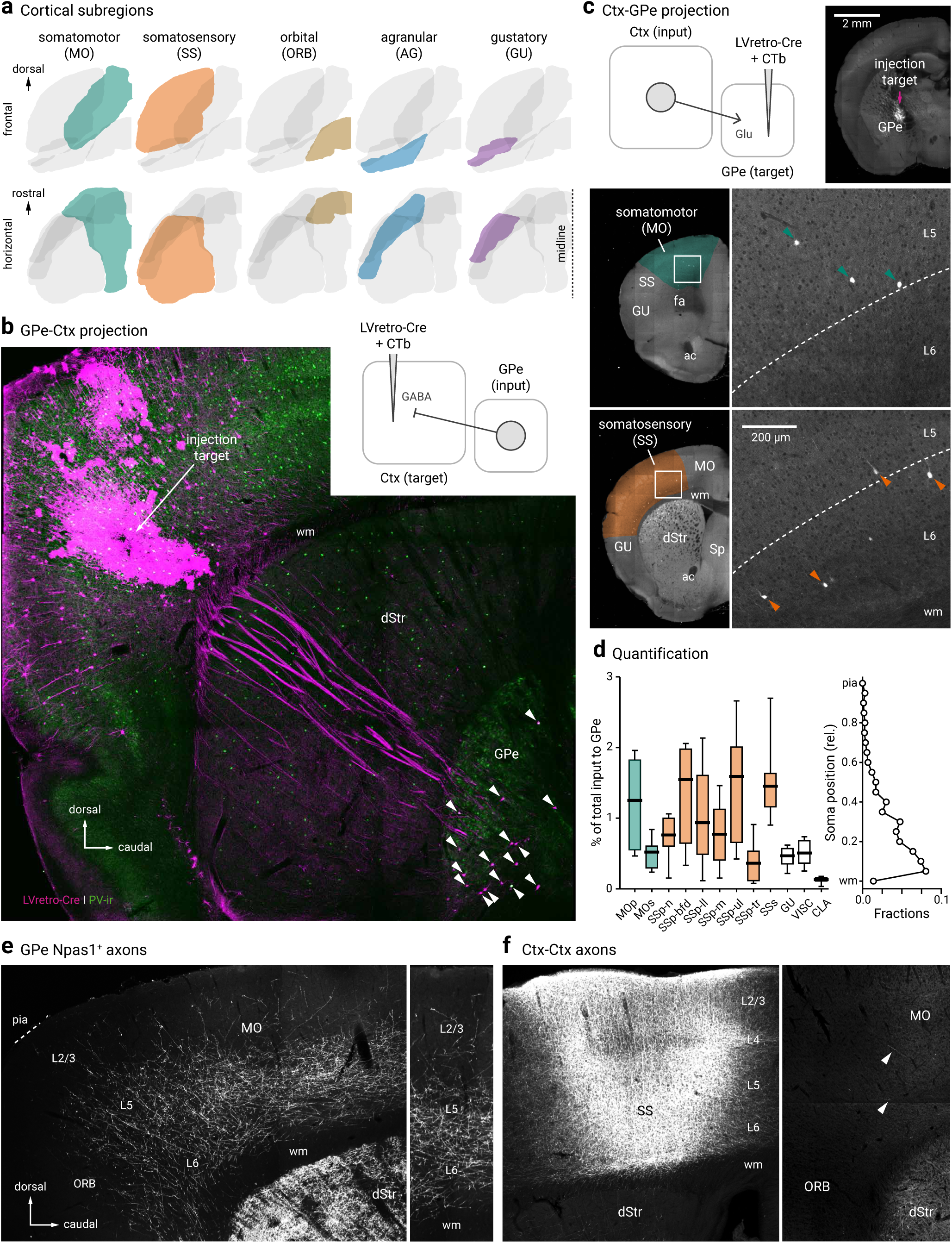
Cortico-pallido-cortical macroscopic anatomy. **a.** Different cortical subregions examined in this study are highlighted. For clarity, frontal (top) and horizontal (bottom) views are shown. **b**. A confocal micrograph showing a representative example of retrogradely-labeled cortex-projecting GPe neurons (arrowhead) using LVretro-Cre in a LSL-tdTomato mouse; PV-immunolabeling (green) was performed in this example. Inset: experimental setup. LVretro-Cre and CTb were injected into different cortical areas mentioned in **a**. **c**. Top left, experimental setup: LVretro-Cre and CTb were injected into the GPe. Cortical inputs to the GPe were mapped using two-photon tomography. Top right, a two-photon image showing the location of the injection site. Bottom, representative two-photon images from coronal sections showing GPe-projecting cortical neurons were found primarily in layer 5 and 6 of MO and SS. **d**. Left, quantification of GPe-projecting neurons across the entire cortex. Medians, interquartile ranges, and 10^th^ to 90^th^ percentiles are represented in a graphical format. Right, laminar position of GPe-projecting neurons. **e**. Low magnification image of Npas1^+^ pallido-cortical axons spanning across ORB, MO, and SS. Note the highest density occurred in layers 5 and 6 of MO followed by SS and dropped off precipitately rostrally in the ORB. Axons extend as far as layer 2/3. **f**. Local cortical infection in a Npas1-Cre-tdTom mouse confirms axons visible in rostral cortical regions are from GPe projection and not ectopic infection of cortical neurons in the caudal areas. Injection site in SS (left) resulted in very low density of caudal to rostral cortico-cortical connectivity in MO and ORB (right). Arrowheads indicate the presence of cortical axons that arose from the more caudal regions. Abbreviations: dStr = dorsal striatum; CLA = claustrum; MOp = primary somatomotor; MOs = secondary somatomotor; ORB = orbital; Sp = septum; SSp-n = primary somatosensory, nose; SSp-bfd = primary somatosensory, barrel field; SSp-ll = primary somatosensory, lower limb; SSp-m = primary somatosensory, mouth; SSp-ul = primary somatosensory, upper limb; SSp-tr = primary somatosensory, trunk; SSs = secondary somatosensory; VISC = visceral; ac = anterior commissure; fa = anterior forceps; wm = white matter.

Though cortical input is known to reach the GPe through the cortico-dStr-GPe and the cortico-STN-GPe pathways (Iwamuro et al., 2017; Jaeger and Kita, 2011; Kita, 2007; Nambu et al., 2000), there is increasing evidence that cortical input targets the GPe directly (Karube et al., 2019; Milardi et al., 2015; Naito and Kita, 1994; Smith and Wichmann, 2015). Using rabies virus tracing, we have recently confirmed the existence of cortical input to GPe neurons (Hunt et al., 2018; Jeong et al., 2016). However, we have not provided a complete representation of the input regions and their relative size. In this study, we sought to map the input from the entire cortical mantle with LVretro-Cre in LSL-tdTomato mice. Automated, serial two-photon tomography was used to provide an unbiased quantification and atlas registration (see Materials and Methods). As expected, input neurons (i.e., tdTomato^+^) from a wide array of brain regions to the GPe were evident (data not shown). A full description of the brain-wide input to the GPe will be detailed in a later publication. The cortical input amounts to ∼10% of the total input to the GPe (*n* = 4,205 out of 45,223 neurons, 8 mice). Consistent with our previous observation, a notable input comes from the SSp followed by MOp. Additionally, but to a much lesser extent, input cells were found primarily in layers 5 and 6 of MOs, SSs, and lateral regions, such as gustatory area, visceral area, and claustrum. These results are summarized in Figure 4c and d.

### Npas1^+^-Nkx2.1^+^ neurons are cortex-projecting

A substantial amount of the cortex-projecting neurons displayed a distinctive larger soma and were located in the caudoventral regions of the GPe—features that are characteristic of ChAT^+^ neurons (Figure 5a). Immunolabeling for ChAT revealed 50% (50 ± 0%, *n* = 19 out of 42 neurons, 3 sections) of tdTomato^+^ neurons were ChAT^+^. The remaining cortex-projecting GPe neurons were ChAT^−^, i.e., non-cholinergic. Our results are highly consistent with prior observations (Ahrlund-Richter et al., 2019; Saunders et al., 2015). However, as the identity of non-cholinergic, cortically-projecting neurons remains elusive, we sought to characterize the molecular profile of these neurons. Through immunolabeling, we identified the ChAT^−^ neurons to be Nkx2.1^+^ (32 ± 6%, *n* = 20 out of 68 neurons, 6 sections) and Npas1^+^ (25 ± 6%, *n* = 41 out of 147 neurons, 13 sections). Furthermore, a population of cortex-projecting neurons was found to be both Nkx2.1^+^ and Npas1^+^ (20 ± 7%, *n =* 8 out of 36 neurons, 3 sections) (Table 4). As expected, the same neurons that expressed Npas1 and Nkx2.1 did not express Foxp2 (0 out of 36 neurons, 3 sections). Similarly, we observed a very small fraction of neurons that were immunoreactive for PV (0 ± 0%, *n =* 3 out of 65 neurons, 8 sections). While abundant retrogradely-labeled neurons were found with injections targeting MOp, SSp, AG and ORB, we observed low levels or non-detectable retrograde labeling with injection to neighboring frontal regions such as secondary somatomotor (MOs, aka M2) and secondary somatosensory (SSs, aka S2) cortices.

**Figure 5.**
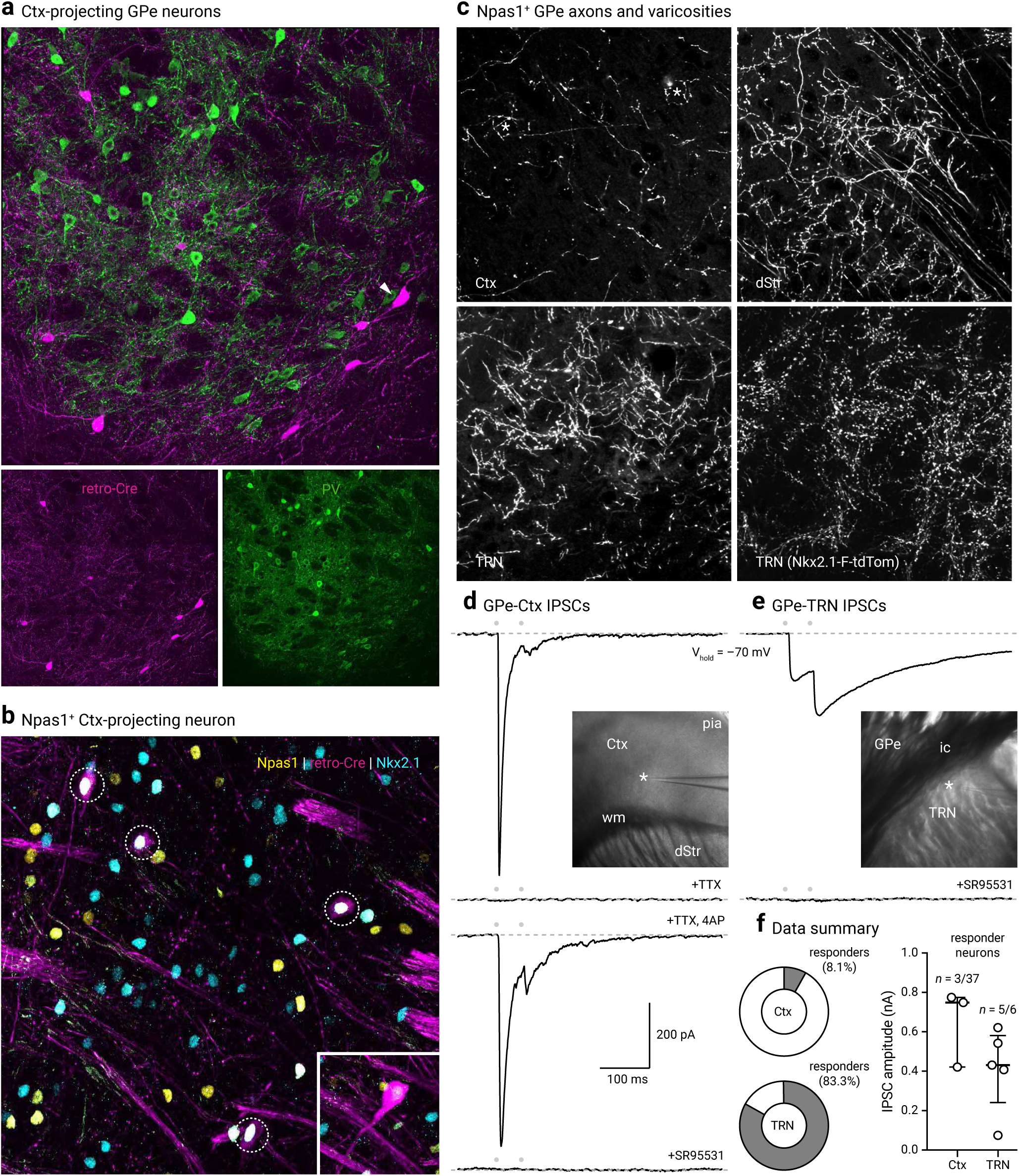
Cortex-projecting neuron properties. **a.** LV retrograde labeled (magenta) GPe neurons with PV immunostaining (green). Note, cortical projecting GPe neurons are not PV^+^. Arrowhead indicates a LV labeled neuron with a large cell body characteristic of cholinergic neurons. **b**. A confocal micrograph showing the co-expression (dotted circles) of Npas1 (yellow) and Nkx2.1 (blue) in cortex-projecting GPe neurons (magenta). Inset: an example of a neuron (shown at the same magnification) that has a large cell body and low Nkx2.1 expression, features of cholinergic neurons within the confines of the GPe. **c**. High magnification confocal micrographs of axons in the Ctx, dStr, and TRN with injection of a CreOn-ChR2 AAV into the GPe of a Npas1-Cre-tdTom mouse. Asterisks in the top left denote putative terminals. Bottom right, high density of synaptic boutons in the TRN of Nkx2.1-F-tdTom mice. **d**. Voltage-clamp recordings of the Npas1^+^ input in a cortical neuron within layers 5 and 6. The recorded neuron was held at –70 mV with a high Cl^−^ electrode; inhibitory postsynaptic currents (IPSCs) were evoked from 20 Hz paired-pulse blue light stimulation (indicated by gray circles). Note the fast and depressing responses evoked. Inset: location of the recorded neuron (asterisk) in the Ctx is shown. IPSCs were attenuated with extracortical stimulation (not shown) and abolished with tetrodotoxin (TTX, 1 µM). Application of 4-aminopyridine (4-AP, 100 µM) in the presence of TTX restored the response with intracortical stimulation. IPSCs were completely blocked with SR95531 (10 µM). **e**. Voltage-clamp recording of a TRN neuron with identical experimental setup shown in **d**. Note the facilitating responses evoked. Inset: location of the recorded neuron (asterisk) is shown. Responses were sensitive to the application of SR95531 (10 µM). **f**. Left, pie charts summarizing the percentages of responders in Ctx and TRN. Right, medians and interquartile ranges of IPSC amplitudes are represented in a graphical format. Abbreviations: dStr = dorsal striatum; Ctx = cortex; TRN = thalamic reticular nucleus; ic = internal capsule; wm = white matter.

To further demonstrate that Npas1^+^ neurons project to the cortex, we injected Npas1-Cre-tdTom mice with a Cre-inducible ChR2-eYFP AAV (see Materials and Methods) into the GPe to trace the arborization patterns of Npas1^+^ axons. Consistent with our previous studies (Glajch et al., 2016; Hernandez et al., 2015), dense Npas1^+^ axons were visible in the dStr. Moreover, we observed axons throughout the MO and SS, but its density dropped off precipitously once passed the ORB. This axonal projection pattern is distinct from that of ChAT^+^ neurons, which arborizes more broadly, including in more caudal regions of the cortex (Moriizumi and Hattori, 1992; Parent et al., 1981; Saunders et al., 2015). To confirm axons observed in the frontal cortex were not a result of ectopic infection of caudal cortical regions, we injected the same Cre-inducible ChR2-eYFP AAV into the SSp region directly above the GPe. We observed only sparse cortico-cortical axons running rostrocaudally (Figure 4f), further confirming that the axons observed in the frontal regions were indeed projections from Npas1^+^ neurons.

The Npas1^+^ axons appeared to arborize heavily in layers 5 and 6 and were present as superficial as layer 2/3 (Figure 4e). Under high-magnification, perisomatic basket-like structures can be found in layer 5 of the MO. In addition, Npas1^+^ axons were found in the TRN, with more moderate projections in the zona incerta (ZI) and SN. Within the SN, we observed a higher density of fibers in the SNc (not shown). The presence of Npas1^+^ axons in the TRN is consistent with the high density of Nkx2.1^+^ synaptic boutons in the area (Figure 5c). As TRN neurons do not express Nkx2.1, the synaptic boutons observed arose from an extrinsic source. These observations suggest Npas1^+^-Nkx2.1^+^ GPe neurons form this projection, and are consistent with previous tracing data that demonstrate an anatomical connection from the GPe to the TRN (Asanuma, 1989, 1994; Clemente-Perez et al., 2017; Cornwall et al., 1990; Gandia et al., 1993; Hazrati and Parent, 1991; Kayahara and Nakano, 1998; Pazo et al., 2013; Shammah-Lagnado et al., 1996).

To evaluate the contact probability and the physiological properties of the synaptic connections from Npas1^+^-Nkx2.1^+^ neurons to the cortex and TRN, Npas1^+^ neurons in the GPe were infected with a ChR2-eYFP AAV; optogenetic-based mapping and patch-clamp recordings were performed from unidentified neurons in the cortex and TRN. As Npas1 axons primarily arborized in cortical layers 5 and 6, neurons in these layers were targeted for recording. In 3 out of 37 cortical neurons recorded, large (438.5–821.7 pA, *n* = 3 neurons) inhibitory postsynaptic currents (IPSCs) were evoked by optogenetic activation of Npas1^+^ axons from the GPe (see Materials and Methods). These were not conducting events, as they were not abolished by the co-application of tetrodotoxin (1 µM) and 4-aminopyridine (100 µM). To confirm the GABAergic nature of the events, SR95531 (a GABA_A_ receptor antagonist, 10 µM) was applied, which completely abolished evoked IPSCs (Figure 5d). In contrast to the apparent low connection probability in the cortex, large IPSCs were readily evoked optogenetically in all but one of the TRN neurons tested (707.8 ± 383.8 pA, *n* = 5 neurons) (Figure 5f). Importantly, as identical optogenetic conditions were used for both experiments, these data argue that the low connection probability detected in the cortex is reflective of selective targeting of cortical neuron subtypes by Npas1^+^ axons. No photocurrents were observed in any of the recorded neurons, ruling out potential contributions of the synaptic events from ectopic infection of neurons local to the recorded areas.

### Pyramidal tract-type cortical neurons target the GPe

Consistent with our two-photon tomography analysis (Figure 4c and d), single-axon reconstruction data from the MouseLight project (Economo et al., 2016) suggest that GPe-projecting cortical neurons are predominantly pyramidal tract (PT)-type. They are thick tufted pyramidal neurons that are located in the lower layer 5 (L5B), do not have cross-hemispheric collaterals, and are multi-projectional (Harris and Shepherd, 2015; Hooks et al., 2018; Kawaguchi, 2017; Kita and Kita, 2012; Shibata et al., 2018) (Figure 6a). This inference corroborated the anterograde viral-tracing data derived from a Sim1-Cre (PT-specific driver line) (Gerfen et al., 2013)—both single-axon (Figure 6a) and bulk tracing data argue that cortical axons enter the GPe (Figure 6c and e). In contrast, similar experiments with Tlx3-Cre (an intratelencephalic-specific driver line) suggest that intratelencephalic neurons do not provide input to the GPe (Figure 6d). Overall, there were general similarities between the topographical organization of the cortico-striatal projections and the cortico-pallidal projections; both frontal and motor projections targeted more rostral areas of the GPe while sensory projections targeted more posterior ones (Hintiryan et al., 2016; Hooks et al., 2018).

**Figure 6.**
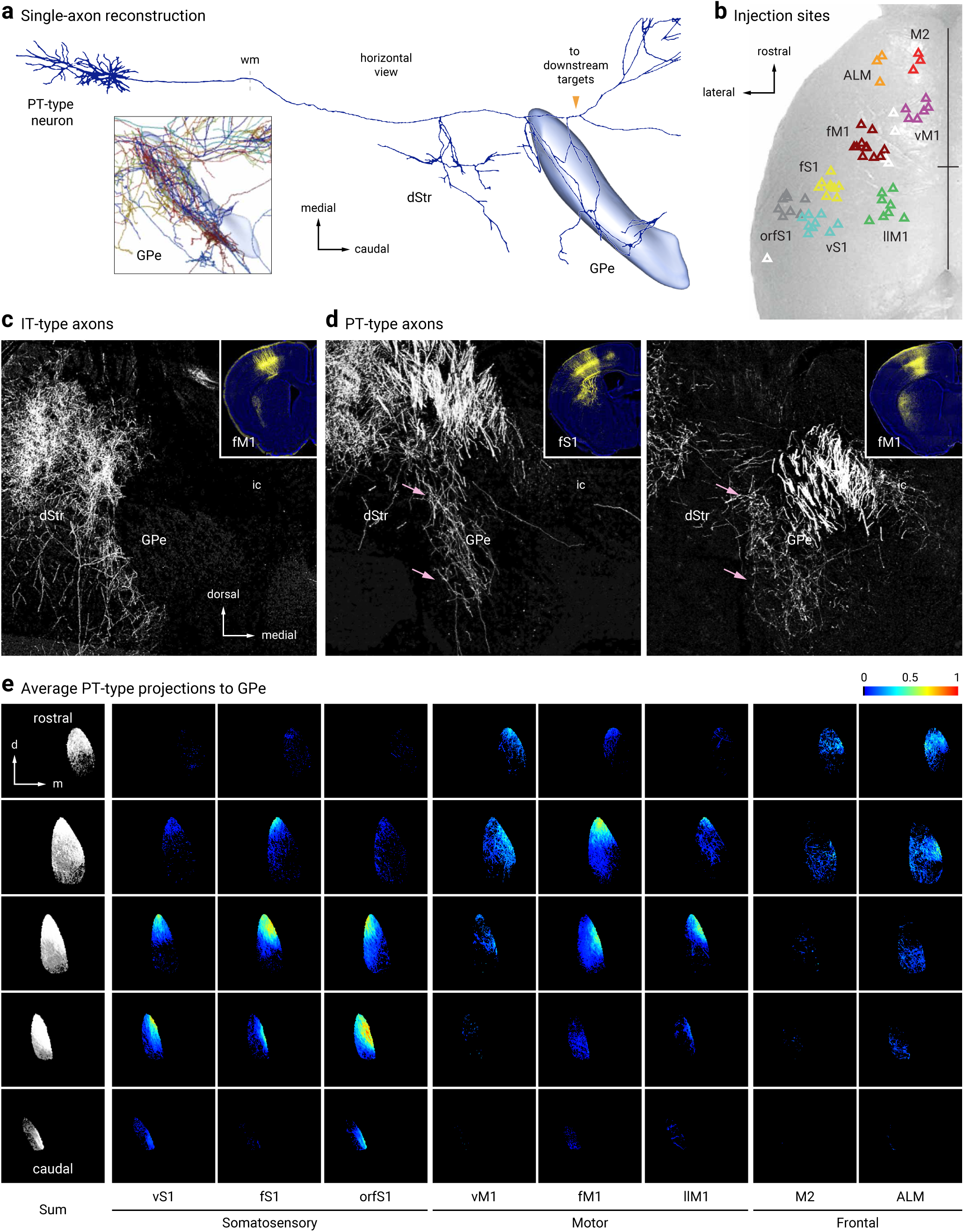
Pyramidal-tract but not intratelencephalic axons collateralize in the GPe. **a.** Single-axon reconstruction of a layer 5 (L5) cortico-pallidal neuron (neuron #AA0122) in the motor cortex. Data are adapted from the MouseLight project (http://mouselight.janelia.org). The axonal projection pattern is consistent with a pyramidal-tract (PT)-type neuron. Inset: axonal arborization of ten different cortical neurons. GPe and axonal endpoint were used as the target location and structure queries, respectively. **b**. Injection site center of mass of Sim1-Cre (L5-PT) plotted and spatially clustered (*n* = 62, triangles). These injection sites correspond to vibrissal, forelimb, and orofacial somatosensory cortices (vS1, fS1, and orfS1); vibrissal, forelimb, and lower limb motor cortices (vM1, fM1, and llM1); and frontal areas (anterior lateral motor cortex (ALM) and secondary motor cortex (M2)). Eight clusters shown in red (M2), orange (ALM), purple (vM1), burgundy (fM1), green (llM1), yellow (fS1), teal (vS1), and gray (orfS1). Indeterminate injection sites are white. Sites are superimposed on an image of the dorsal surface of mouse cortex. Black cross marks indicate midline and bregma. For simplicity, injection sites in Tlx3-Cre (L5-IT) are not shown (see Hooks et al., 2018, for further information). **c–d**. Tlx3-Cre (IT-type) projections from fM1 in dStr but not GPe. Sim1-Cre (PT-type) projections from fS1 (left) and fM1 (right) in dStr and GPe (pink arrows). Inset: Coronal images of injection sites in Tlx3-Cre and Sim1-Cre showing the cell body locations and their axonal projections. **e.** Coronal images of the average normalized PT-type projection to GPe from eight cortical areas. Each column is a cortical projection, with rows going from anterior (top) to posterior (bottom). Each projection is normalized for comparison within the projection.

To provide a confirmation of the existence of synaptic terminals formed by cortical axons in the GPe, we injected AAVretro-ChR2-eYFP into the GPe of Emx1-Cre mice (Gorski et al., 2002). This resulted in selective ChR2-eYFP expression in cortical neurons and their axons in the GPe, STN, and dStr (Figure 7a). A high density of labeled neurons (and processes) were observed within the ORB, MO, and SS. This expression pattern is in agreement with our two-photon tomography data (Figure 4c and d) and the axonal projection patterns revealed by Sim1-Cre (Figure 6c and e). Furthermore, these data reinforce the idea that the cortical axons observed in the GPe in both the single axon and bulk tracking studies were not simply passage fibers. In keeping with this idea, co-localization of VGluT1 in ChR2-eYFP-labeled varicosities was readily observed (Figure 7b). To provide more definitive evidence, *ex vivo* voltage-clamp recordings were performed; optogenetic activation of cortical input evoked EPSCs in 20 out of 25 GPe neurons. However, the size of the EPSCs (29.0 ± 18.2 pA) spans over three orders of magnitude in the recorded neurons. Consistent with the single-axon and bulk tracing data, which both show that cortico-pallidal neurons produce collaterals within the dStr, EPSCs (100.8 ± 79.1 pA) were evoked readily across all striatal projection neurons tested (*n* = 6) (Figure 7d).

**Figure 7.**
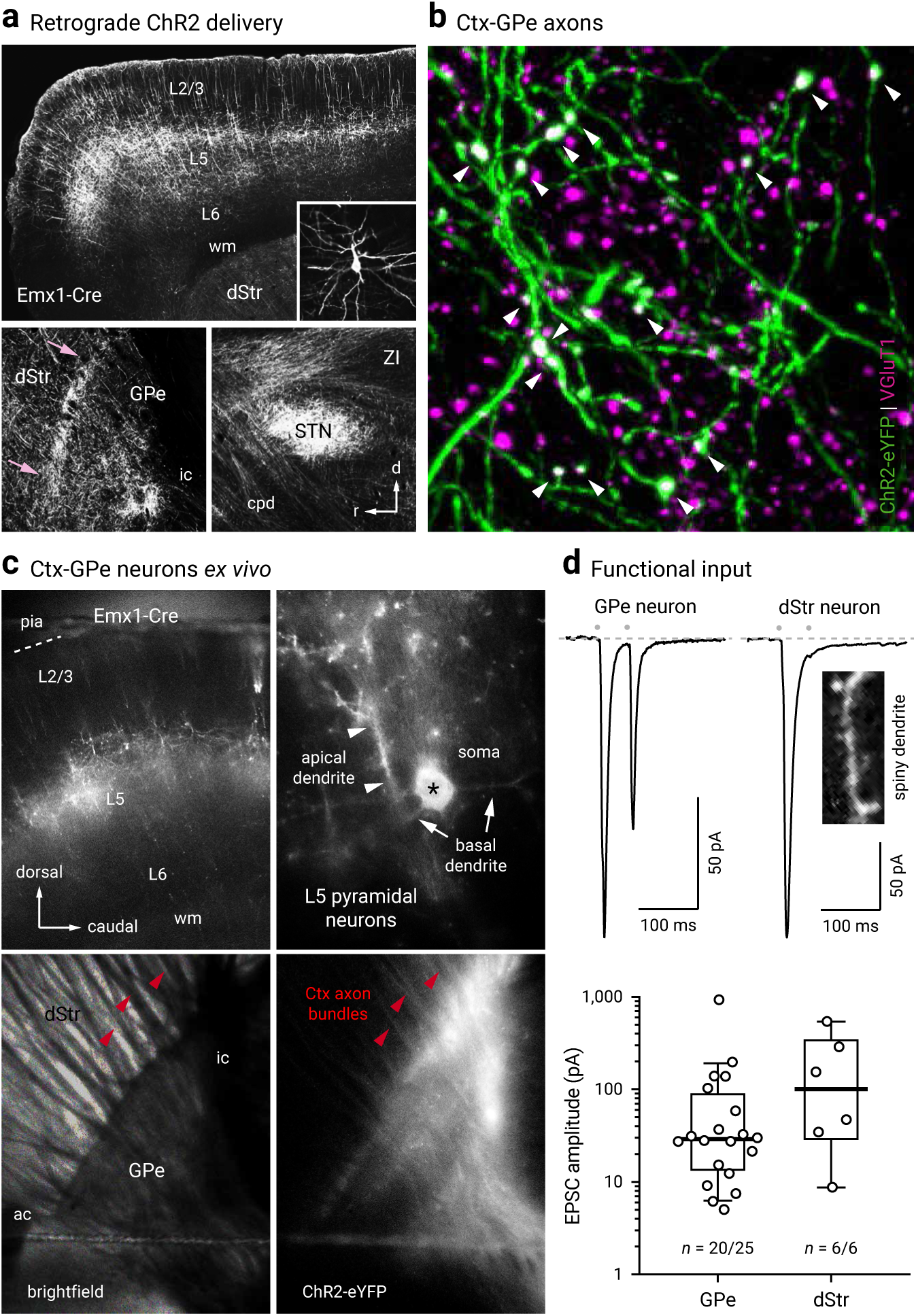
Cortex neurons form functional synapses on GPe neurons. **a.** Top, a confocal micrograph of a sagittal section from an Emx1-Cre mouse, showing the neuronal elements expressing ChR2-eYFP delivered from an AAVretro injection into the GPe. Inset, a ChR2-eYFP^+^ layer 5 neuron with morphology typical of pyramidal neurons is shown. Bottom left, cortical axons was observed at the GPe level. An enrichment of axons was present at the rostral pole of the GPe, immediate adjacent to the dStr (pink arrows). Bottom right, a high density of cortico-pallidal axons were observed to collateralize in the subthalamic nucleus (STN). **b**. A high magnification image showing the co-localization (white arrowheads) of VGluT1 (magenta) in cortico-pallidal axons (green). **c**. Epifluorescence image from *ex vivo* tissue showing robust expression of ChR2-eYFP in cortical neurons. Top, AAVretro-CreOn-ChR2-eYFP was injected into the GPe of an Emx1-Cre mouse. Robust ChR2-eYFP expression in the MO was observed (left). ChR2-eYFP^+^ neurons were readily seen at high magnification (right). Asterisk indicates the soma of a typical ChR2-eYFP^+^ neuron. Bottom, cortico-pallidal axons were preserved in an *ex vivo* slice preparation. **d**. Top, functional cortical inputs were recorded in GPe neurons (20 out of 25) and dStr projection neurons (6 out of 6). EPSCs were evoked with optogenetics. Bottom, box and scatter plots summarizing EPSC amplitude recorded from GPe neurons (left) and dStr SPNs (right). Note the large variance in the data. Red arrowheads: cortical axon bundles. Orange arrowheads: cortical axons projecting to regions caudal to the GPe. White arrowheads: apical dendrites. Abbreviations: dStr = dorsal striatum; GPe = external globus pallidus; STN = subthalamic nucleus; ZI = zona incerta; ac = anterior commissure; cpd = cerebral peduncle; ic = internal capsule; wm = white matter.

### Lhx6^+^ neuron subtypes have unique spatial patterns

Spatial distribution may vary with neuronal identity, as seen in the cortex, for example, where neurons are organized in a highly laminar-specific manner (Harris and Shepherd, 2015; Huang et al., 2007; Wamsley and Fishell, 2017). We therefore integrated spatial distribution as a metric to phenotype different GPe neuron types (Figure 8 and 9). Overall, more neurons populate towards the rostral pole of the GPe. We noted that ChAT^+^ neurons and Lhx6^+^-Sox6^+^ neurons are displaced more heavily toward the caudoventral regions and rostroventral regions, respectively. All other identified neurons are more evenly distributed throughout the GPe. This analysis, however, does not capture any lateromedial gradients. As neurons were routinely sampled from three different lateromedial planes across the GPe, the abundance of identified neurons were tallied. Consistent with previous studies (Hernandez et al., 2015; Mastro et al., 2014), we observed a lateromedial gradient of PV^+^ neurons (lateral = 46 ± 10%, *n* = 4,368 neurons, 24 sections; intermediate = 45 ± 11%, *n* = 5,113 neurons, 27 sections; medial = 32 ± 7%, *n* = 2,829 neurons, 20 sections) and Lhx6^+^ neurons (lateral = 28 ± 6%, *n* = 1,422 neurons, 10 sections; intermediate = 42 ± 9%, *n* = 2,050 neurons, 10 sections; medial = 45 ± 12%, *n* = 2,190 neurons, 8 sections). PV^+^ neurons were more concentrated in the lateral than the medial level; the reverse pattern was found for Lhx6^+^ neurons. In Figure 9, we illustrate the distribution of Lhx6^+^-Sox6^−^ neurons (lateral = 8 ± 4%, *n* = 187 neurons, 5 sections; intermediate = 10 ± 1%, *n* = 292 neurons, 5 sections; medial = 21 ± 7%, *n* = 459 neurons, 5 sections), which follow the same pattern as pan-Lhx6^+^ neurons. While PV^+^-Lhx6^+^ neurons displayed a similar pattern (lateral = 8 ± 2%, *n* = 116 neurons, 4 sections; intermediate = 11 ± 6%, *n* = 202 neurons, 3 sections; medial = 8 ± 1%, *n* = 175 neurons, 3 sections) as pan-Lhx6^+^ neurons, Npas1^+^-Lhx6^+^ neurons do not (lateral = 14 ± 2%, *n* = 231 neurons, 4 sections; intermediate = 15 ± 2%, *n* = 308 neurons, 4 sections; medial = 14 ± 2%, *n* = 284 neurons, 4 sections). The lateromedial gradients of different GPe neuron types are summarized in Figure 9.

**Figure 8.**
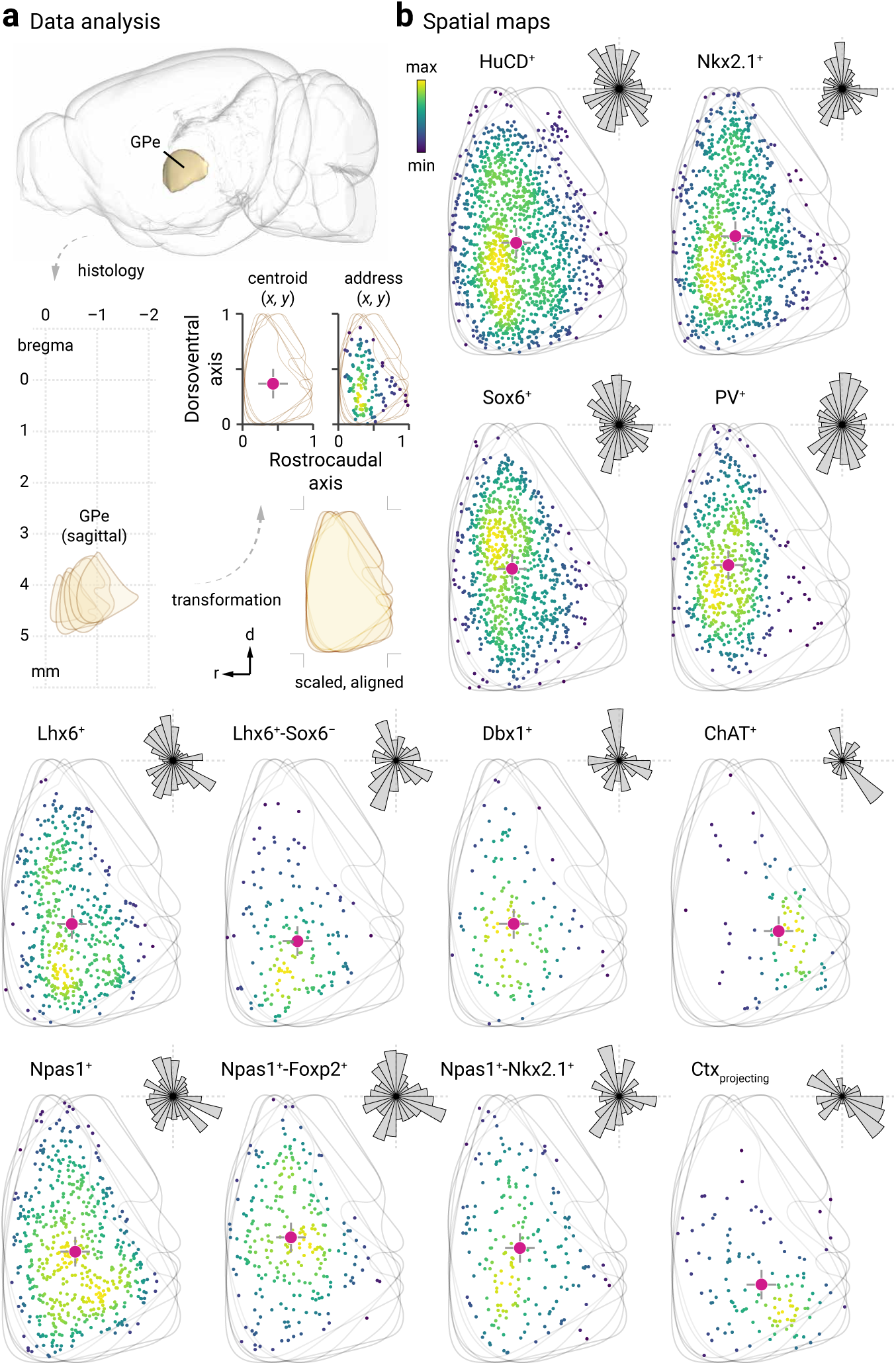
Lateromedial gradients and relative abundance of different GPe neuron classes. **a.** Spatial maps of the pan-Lhx6^+^ and unique Lhx6^+^ Sox6^−^ GPe neuron populations. Both populations display a lateromedial gradient with more neurons populating the medial GPe. **b**. Relative abundance of neuron classes in different lateromedial subdivisions of the GPe (Sox6^+^ = lateral: 60 ± 12%, *n* = 4,457 neurons, intermediate: 63 ± 11%, *n* = 5,112 neurons, medial: 50 ± 11%, *n* = 3,286 neurons; Nkx2.1^+^ = lateral: 56 ± 7%, *n* = 3,365 neurons, intermediate: 53 ± 9%, *n* = 3,878 neurons, medial: 64 ± 14%, *n* = 3,265 neurons; PV^+^ = lateral: 46 ± 10%, *n* = 4,368 neurons, intermediate: 45 ± 11%, *n* = 5,113 neurons, medial: 32 ± 7%, *n* = 2,829 neurons; Lhx6^+^ = lateral: 28 ± 6%, *n* = 1,422 neurons, intermediate: 42 ± 9%, *n* = 2,050 neurons, medial: 45 ± 12%, *n* = 2,190 neurons; Npas1^+^ = lateral: 32 ± 6%, *n* = 2,635 neurons, intermediate: 31 ± 6%, *n* = 2,903 neurons, medial: 27 ± 7%, *n* = 2,252 neurons; Foxp2^+^ = lateral: 24 ± 3%, *n* = 939 neurons, intermediate: 26 ± 4%, *n* = 1,115 neurons, medial: 25 ± 6%, *n* = 686 neurons; Dbx1^+^ = lateral: 10 ± 2%, *n* = 1,219 neurons, intermediate: 9 ± 2%, *n* = 1,540 neurons, medial: 8 ± 2%, *n* = 1,121 neurons; ChAT^+^ = lateral: 6 ± 1%, *n* = 100 neurons, intermediate: 4 ± 1%, *n* = 76 neurons, medial: 6 ± 3%, *n* = 91 neurons). Percentage total was calculated from HuCD^+^ cells within each section. Note that PV and Npas1 were expressed in a largely non-overlapping fashion (2 ± 2%, *n* = 96 neurons, 12 sections). In contrast, considerable overlap between Lhx6 and PV (28 ± 7%, *n* = 654 neurons, 12 sections) or Npas1 (35 ± 5%, *n* = 818 neurons, 12 sections) was observed; the remaining fraction was uniquely labeled with Lhx6. Medians and interquartile ranges are represented in a graphical format. Asterisks denote statistical significance level: \*\**P* < 0.01, Mann–Whitney *U* test.

**Figure 9.**
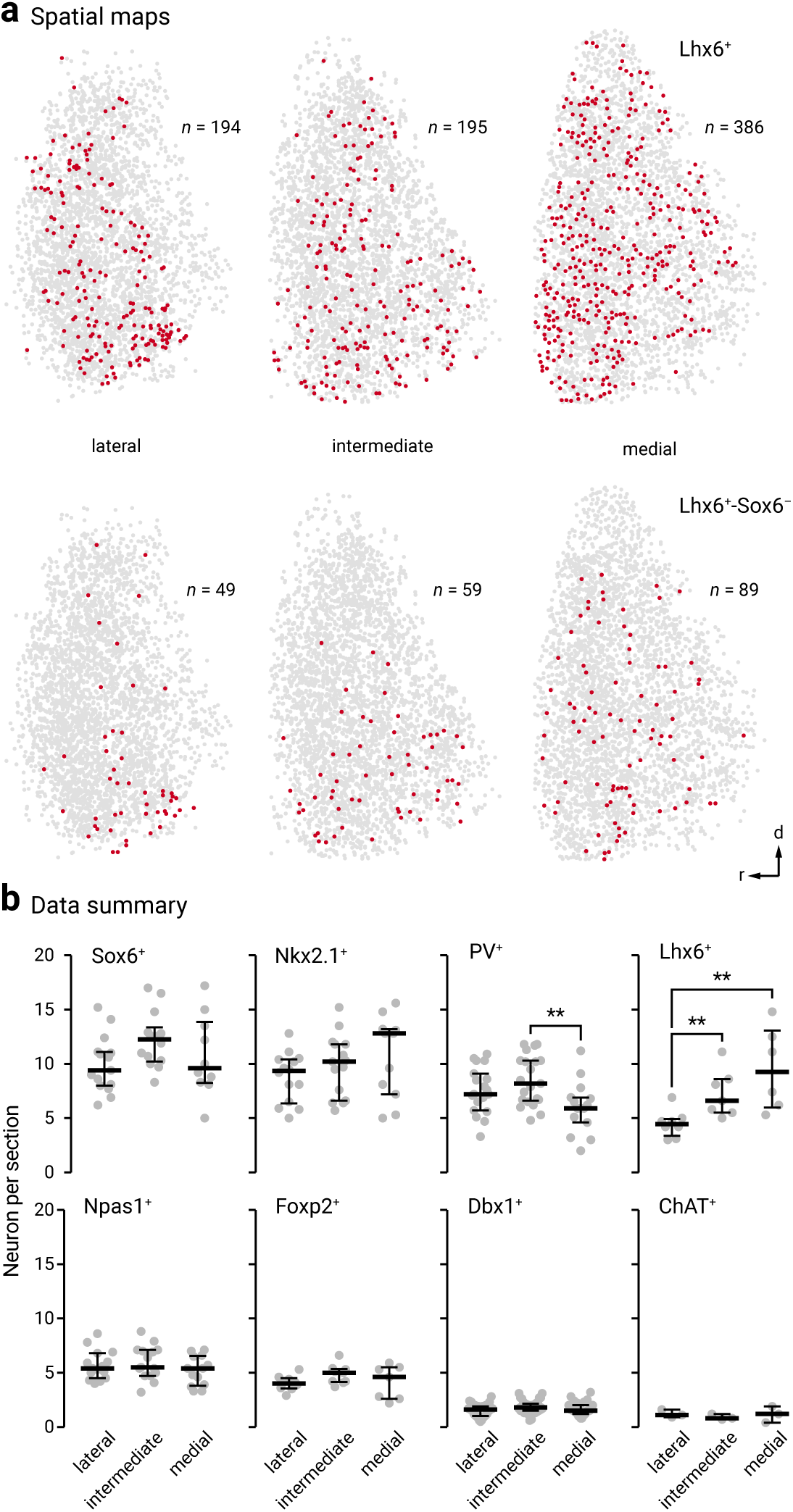
Spatial distribution of GPe neuron subtypes. **a.** Spatial information of GPe neurons cannot be represented relative to bregma location (top, lower left) because of its complex geometry. To mathematically describe the spatial distribution of GPe neurons in each brain section, fixed mouse brains were sagittally sectioned and histologically processed. Images were manually aligned to a reference atlas. GPe neurons located at six different lateromedial levels (2.73 mm, 2.53 mm, 2.35 mm, 2.15 mm, 1.95 mm, and 1.73 mm) were charted and collapsed onto a single plane. As the GPe is similarly-shaped across the lateromedial extent (lower right), both the rostrocaudal and dorsoventral extent were assigned to 0 and 1. The address of each neuron is defined by their x-y coordinates and represented with a marker (black). To capture the aggregate spatial distribution, a geometric centroid (red) of each neuron population was then determined to represent the center of mass in both x and y dimensions. Centroids are then used as the origin for the polar histograms in **b**. Size of each sector represents the relative neuron count as a function of direction. **b**. Representative data of neurons from two individual mice are shown in each case, except for retrogradely-labeled cortically-projecting neurons (*n* = 7 mice; 119 neurons; 15 sections). Each marker represents a neuron. The density of neurons are encoded with a yellow-blue gradient. Hash marks, which represent the dorsoventral and rostrocaudal axes, are presented with the centroids and polar histograms to facilitate comparison. Bin sizes in the polar histograms were chosen based on the size of each neuron population. The (x, y) centroid values for the respective GPe distributions were: HuCD^+^ (0.3798, 0.4168); Nkx2.1^+^ (0.3599, 0.4439); Sox6^+^ (0.3587, 0.4529); PV^+^ (0.3205, 0.4699); Lhx6^+^ (0.3918, 0.3827); Lhx6^+^-Sox6^−^ (0.3755, 0.3164), Dbx1^+^ (0.3679, 0.3828); ChAT^+^ (0.6024, 0.3569); Npas1^+^ (0.4106, 0.4140), Npas1^+^-Foxp2^+^ 0.3695, 0.4676); Npas1^+^-Nkx2.1^+^ (0.4026, 0.4278); Ctx-projecting GPe neurons (0.5061, 0.2911).

### GPe neuron subtypes have distinct intrinsic properties

To further define GPe neuron subtypes, *ex vivo* electrophysiological analyses were performed systematically on genetically-identified GPe neuron subtypes, including the less well-studied neuron subtypes. We used recording and analysis routines (see Materials and Methods) identical to those used in our previous study to facilitate cross-comparison between the two (Hernandez et al., 2015).

To identify Foxp2^+^ neurons, we infected Foxp2-Cre mice in the GPe with a CreOn-mCherry AAV (see Materials and Methods). To confirm the validity of the approach, a subset of these mice were examined for cellular specificity of Cre-mediated mCherry expression (mCherry^+^). In nearly all mCherry^+^ neurons examined, Foxp2 was expressed (100 ± 0%, *n* = 473 out of 485 neurons, 6 sections). No GPe neurons expressed mCherry when the same virus was injected in wild-type mice (*n* = 2 mice, 6 sections). For electrophysiological analysis, we qualitatively categorized Lhx6^+^ neurons as “bright” and “dim” based on their GFP expression level (Figure 10a), though the definitive identities of Lhx6^+^_bright_ and Lhx6^+^_dim_ neurons were not confirmed *post hoc*. To identify PV^+^-Dbx1^+^ neurons, an intersectional cross was made—PV-Dbx1-FL-tdTom (PV-Flp;Dbx1-Cre;LSL-FSF-tdTomato)—to label PV^+^-Dbx1^+^ neurons (tdTomato^+^) (Figure 10a). To unequivocally identify Npas1^+^-Lhx6^+^ neurons, which are equivalent to Npas1^+^-Nkx2.1^+^ neurons, we crossed Npas1-tdTom and Lhx6-GFP mice. Double-positive (tdTomato^+^ and GFP^+^) neurons were targeted for recordings. ChAT^+^ neurons, which can be identified based on their unique somatodendritic morphology, were not included in this analysis as we have previously established that they have a very distinct electrophysiological profile (Hernandez et al., 2015). Lastly, because we do not have genetic access to Lhx6^+^-Sox6^−^ neurons, we did not have a means to target them for direct recording.

**Figure 10.**
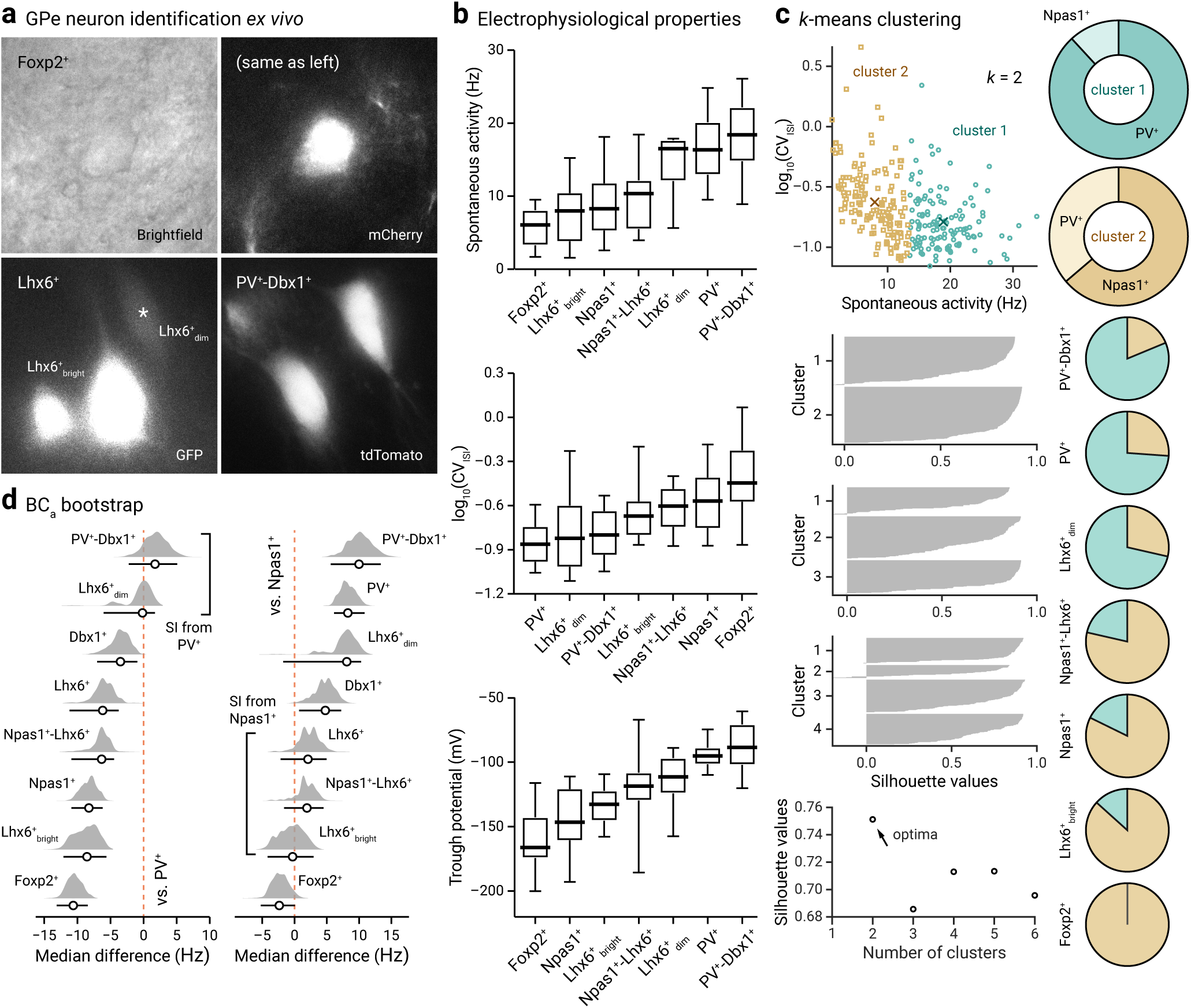
Genetically-identified GPe neurons differ in their spontaneous activity. **a.** Representative brightfield and epifluorescence images of GPe neuron subtypes in *ex vivo* brain slices. Foxp2^+^ neuron (top, brightfield and mCherry), Lhx6^+^_bright_ neurons and Lhx6^+^_dim_ (bottom left, GFP), and PV^+^-Dbx1^+^ neurons (bottom right, tdTomato) were captured at 60x magnification. Note the difference in the morphology and GFP expression among Lhx6^+^ neurons. **b**. Box-plot summary of the electrophysiological properties of identified GPe neuron subtypes. Data are ranked based on the median values. See Table 5 and 6 for median values, sample sizes, and statistical analysis. Medians, interquartile ranges, and 10^th^ to 90^th^ percentiles are represented in a graphical format. **c**. Top left, visualization of the clustered data on spontaneous activity (rate and CV_ISI_) for *k* = 2 clusters. Centroid values for cluster 1 (teal circle) and 2 (tan squares) are: 18.9 Hz, 0.16 and 7.9 Hz, 0.24. Middle left, silhouette plots for different clusters. Bottom right, silhouette values are plotted against cluster numbers showing an optima at *k* = 2. Large positive silhouette values indicate that the data point is close to its cluster’s centroid, whereas negative silhouette values indicate that the data point is closer to the centroid of the other cluster. Right, a series of pie charts showing the membership assignment of different genetically-defined GPe neuron subtypes. The membership assignment in cluster 1 (teal) and 2 (tan) for each neuron subtypes are: PV^+^-Dbx1^+^ (81.3%, 18.8%, *n* = 16), PV^+^ (73.9%, 26.1%, *n* = 111), Lhx6^+^_dim_ (71.4%, 28.6%, *n* = 7), Npas1^+^-Lhx6^+^ (21.4%, 78.6%, *n* = 14), Npas1^+^ (17.7%, 82.3%, *n* = 62), Lhx6^+^_bright_ (13.3%, 86.7%, *n* = 15), Foxp2^+^ (0.0%, 100.0%, *n* = 20). Data are not shown for Dbx1^+^ (50.0%, 50.0%, *n* = 20) and Lhx6^+^ (61.1%, 38.9%, *n* = 18), which both contain a mixture of PV^+^ neurons and Npas1^+^ neurons. **d**. Bias-corrected and accelerated (BC_a_) bootstrap estimation of effect sizes (median differences) and 95% confidence intervals. The median difference in spontaneous rate for seven comparisons against the PV^+^ neurons (left) and Npas1^+^ neurons (right) are shown. Median differences are plotted as bootstrap sampling distributions. Each median difference is depicted as a circle. Median differences are also encoded by color saturation. Lower and upper confidence interval bounds are indicated by the horizontal bars. Lhx6^+^_dim_ neurons and PV^+^-Dbx1^+^ neurons are statistically non-significant from PV^+^ neurons (*P* = 0.452 and 0.244). Lhx6^+^_bright_ neurons, Npas1^+^-Lhx6^+^ neurons, and Lhx6^+^ neurons are statistically non-significant from Npas1^+^ neurons (*P* = 0.440, *P* = 0.292, and 0.066).

**Table 5.**
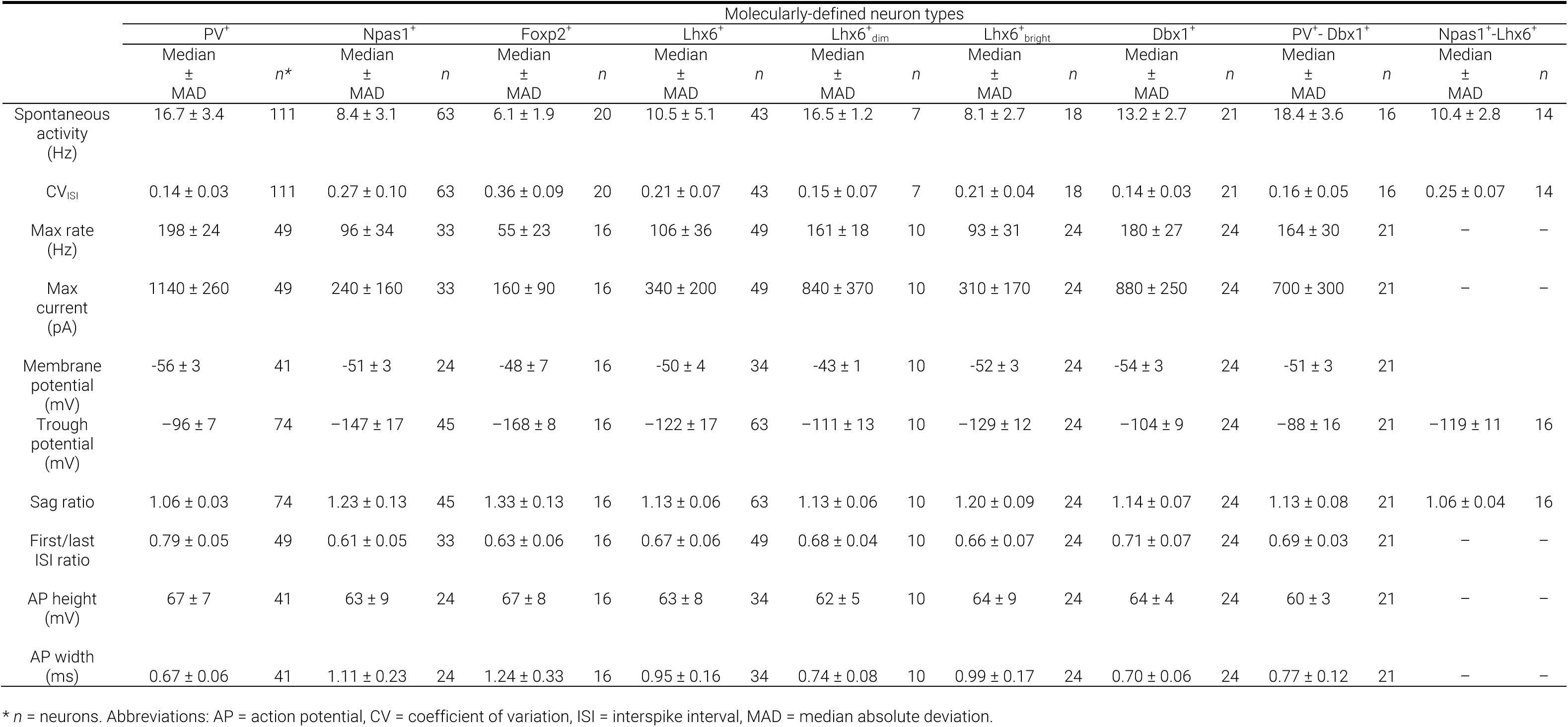
Electrophysiological characteristics of GPe neurons

**Table 6.**
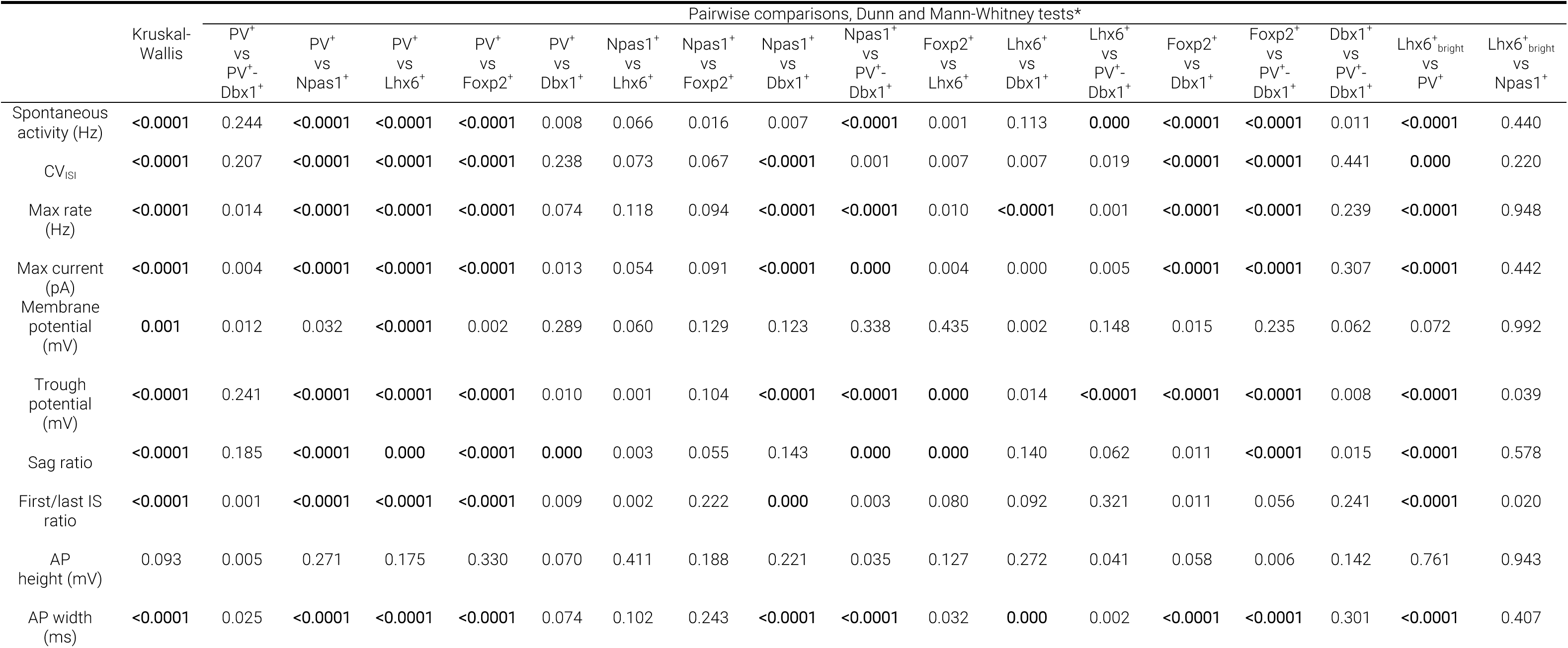
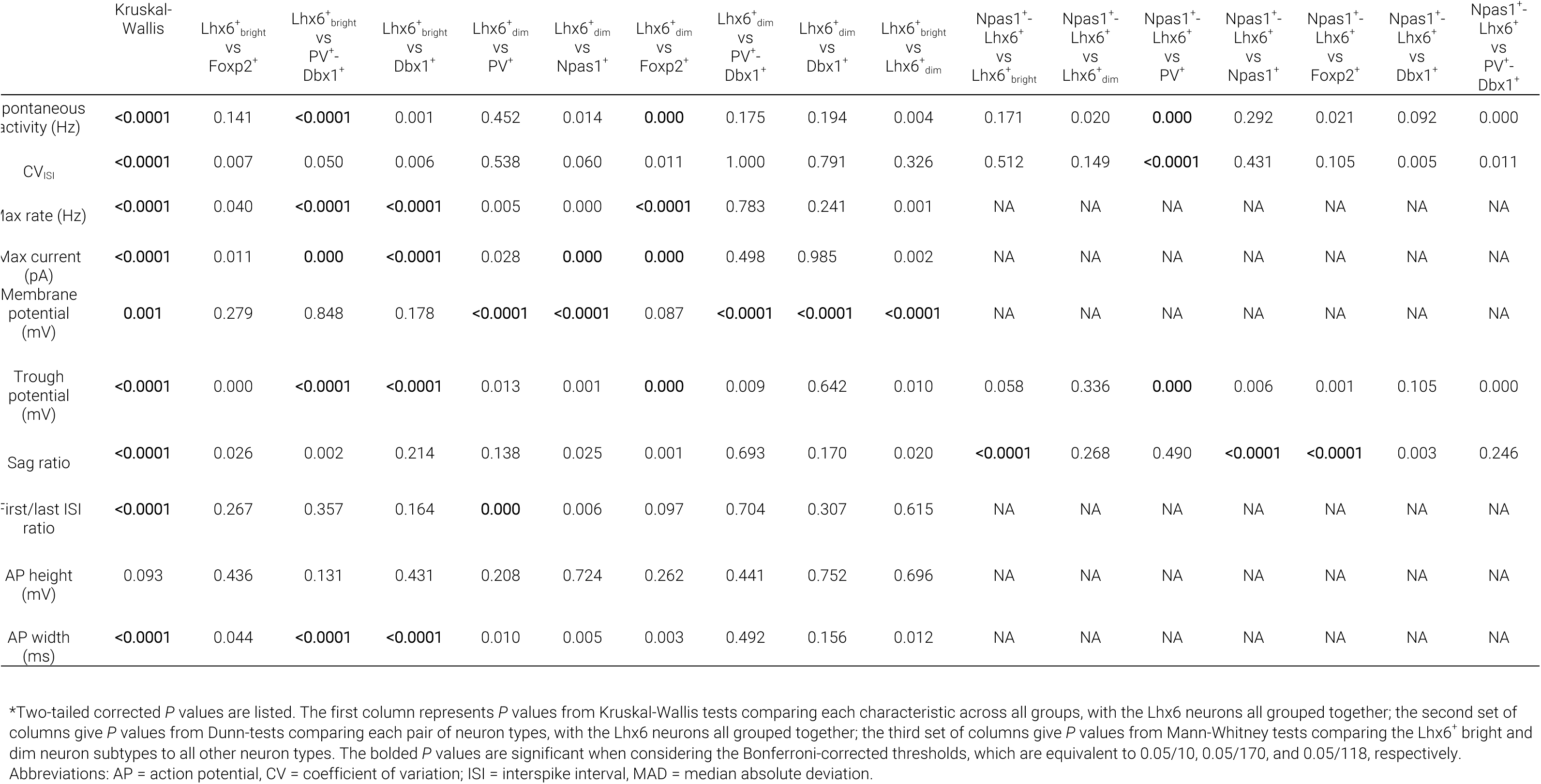
Statistical analysis for electrophysiological characteristics

To highlight, Foxp2^+^ neurons and PV^+^-Dbx1^+^ neurons were at the extremes in terms of their spontaneous activity level measured in cell-attached recordings (Foxp2^+^ = 6.1 ± 1.9 Hz, *n* = 20; PV^+^-Dbx1^+^ = 18.4 ± 3.6 Hz, *n* = 16), with the rest of the neuron subtypes displaying properties traversing the spectrum. Furthermore, our results corroborate findings from our prior studies (Hernandez et al., 2015) that PV^+^ neurons fire at a higher rate than Npas1^+^ neurons (PV^+^ = 16.7 ± 3.4 Hz, *n* = 111; Npas1^+^ = 8.4 ± 3.1 Hz, *n* = 63, *P* < 0.0001). Lhx6^+^ neurons exhibited firing rates that are in between PV^+^ neurons and Npas1^+^ neurons (Lhx6^+^ = 10.5 ± 5.1 Hz, *n* = 43). Within the Lhx6^+^ population, Lhx6^+^_bright_ neurons and Lhx6^+^_dim_ neurons exhibited different firing rates (Lhx6^+^_bright_= 8.1 ± 2.7 Hz, *n* = 18; Lhx6^+^_dim_= 16.5 ± 1.2 Hz, *n* = 7, *P* = 0.004). Additionally, the Npas1^+^-Lhx6^+^ neurons had a higher spontaneous firing than the Foxp2^+^ population (Npas1^+^-Lhx6^+^ = 10.4 ± 2.8 Hz, *n* = 14, *P* = 0.021) but was comparable to Lhx6^+^_bright_ neurons (*P* = 0.171) (Figure 10b). As expected, Dbx1^+^ neurons exhibited firing behavior that was most consistent with its composition, with most neurons exhibiting similar firing rates to PV^+^ neurons and some more similar to Npas1^+^ neurons (See Table 5 and 6, Dbx1^+^ = 13.2 ± 2.7 Hz, *n* = 21).

As we have previously shown that the regularity of the firing varied with GPe neuron types (Hernandez et al., 2015), we measured the coefficient of variation in the interspike-interval (CV_ISI_). Consistent with our prior observations, PV^+^ neurons exhibited a lower CV_ISI_ than Npas1^+^ neurons (PV^+^ = 0.14 ± 0.03, *n* = 111; Npas1^+^ = 0.27 ± 0.10, *n* = 63, *P* < 0.0001). Dbx1^+^ neurons and PV^+^-Dbx1^+^ neurons had CV_ISI_ that were statistically similar to those of PV^+^ neurons (Dbx1^+^ = 0.14 ± 0.03, *n* = 21; PV^+^-Dbx1^+^ = 0.16 ± 0.05, *n* = 16). Foxp2^+^ neurons had the largest variability in their firing rate, as indicated by the highest CV_ISI_ within the GPe neuronal population (Foxp2^+^ = 0.36 ± 0.09, *n* = 20). Npas1^+^-Lhx6^+^ neurons had CV_ISI_ statistically similar to that of the Lhx6^+^_bright_ neurons (Npas1^+^-Lhx6^+^ = 0.25 ± 0.07, *n* = 14, Lhx6^+^_bright_ = 0.21 ± 0.04 *n* = 18, *P* = 0.512).

To provide comprehensive electrophysiological profiles, we investigated a range of electrophysiological properties of GPe neurons in whole-cell current-clamp recordings. As seen in prior studies, PV^+^ neurons had the highest maximum firing rate (PV^+^ = 198 ± 24 Hz, *n* = 49). Lower maximum firing rates were observed in Npas1^+^ neurons and Foxp2^+^ neurons (Npas1^+^ = 96 ± 34 Hz, *n* = 33; Foxp2^+^ = 55 ± 23, *n* = 16) when compared to PV^+^, Dbx1^+^, and PV^+^-Dbx1^+^ neurons (*P* < 0.0001 for all). Within the Lhx6 population, Lhx6^+^_bright_ neurons and Lhx6^+^_dim_ neurons exhibited different maximum firing rates (Lhx6^+^_bright_= 93 ± 31 Hz, *n* = 24; Lhx6^+^_dim_= 161 ± 18 Hz, *n* = 10, *P* = 0.001). GPe neuron subtypes also exhibited different responses to hyperpolarizing current injections (Figure 10b). More negative trough potentials were noted in the Npas1^+^ neurons and Foxp2^+^ neurons (Npas1^+^ = –147 ± 17 mV, *n* = 45; Foxp2^+^ = –168 ± 8 mV, *n* = 16) when compared to PV^+^ neurons (PV^+^ = –96 ± 7 mV, *n* = 74, *P* < 0.0001 for both). Within Lhx6^+^ neurons, Lhx6^+^_bright_ neurons and Lhx6^+^_dim_ neurons exhibited different trough potentials (Lhx6^+^_bright_= – 129 ± 12 mV, *n* = 24, Lhx6^+^_dim_= –111 ± 13 mV, *n* = 10, *P* = 0.010). The trough potential observed in Npas1^+^-Lhx6^+^ neurons (Npas1^+^-Lhx6^+^ = –119 ± 11 mV, *n* = 16) was less negative than both Npas1^+^ neurons (*P* = 0.006) and Foxp2^+^ neurons (*P* = 0.001). As seen in previous work from our lab (Hernandez et al., 2015), PV^+^ and PV^+^-Dbx1^+^ neurons exhibited the least negative trough potential compared to the other studied GPe neuron types and were not different from one another (*P* = 0.241) (PV^+^ = –96 ± 7 mV, *n* = 74, PV^+^-Dbx1^+^ = –88 ± 16 mV, *n* = 21). Dbx1^+^ neurons had trough potential (Dbx1^+^ = –104 ± 9 mV, *n* = 24) more negative than those seen in PV^+^ neurons (*P* = 0.010) and PV^+^-Dbx1^+^ neurons (*P* = 0.008). Similar to the trough potentials, higher sag ratios were observed for Npas1^+^ neurons and Foxp2^+^ neurons (Npas1^+^ = 1.23 ± 0.13, *n* = 45; Foxp2^+^ = 1.33 ± 0.13, *n* = 16). PV^+^ neurons had the lowest sag ratio (PV^+^ = 1.06 ± 0.03, *n* = 74). Within Lhx6, a difference in the sag ratio was observed between Lhx6^+^_bright_ neurons and Lhx6^+^_dim_ neurons (Lhx6^+^_bright_= 1.20 ± 0.09, *n* = 24, Lhx6^+^_dim_= 1.13 ± 0.06, *n* = 10, *P* = 0.020). The Npas1^+^-Lhx6^+^ neurons exhibited a lower sag ratio than both Npas1^+^ neurons (Npas1^+^-Lhx6^+^ = 1.06 ± 0.04, *n* = 16, *P* < 0.0001) and Foxp2^+^ neurons (*P* < 0.0001). As expected, the PV^+^-Dbx1^+^ neurons had statistically similar sag ratios to the PV^+^ neuron population (PV^+^-Dbx1^+^ = 1.13 ± 0.08, *n* = 21, *P* = 0.185). A full description of the electrophysiological characteristics of GPe neurons is listed in table form (Table 5).

As the spontaneous activity level and regularity vary with neuron types, we implemented *k*-means analysis to examine if GPe neurons can be classified into electrophysiological subtypes (Figure 10c). This analysis returned an optima of two clusters with the centroids at (7.9 Hz, 0.24; *n* = 297 and 18.9 Hz, 0.16; *n* = 259), which approximate to the median values of Npas1^+^ neurons (8.4 Hz, 0.27) and PV^+^ neurons (16.7 Hz, 0.14), respectively. To evaluate the molecular correspondence of these two mathematically-defined clusters composition, their compositions were visualized. Cluster 1 primarily consists of PV^+^ neurons (88.2%) and cluster 2 Npas1^+^ neurons (63.8%).

The *k*-means clustering further supports the notion that PV^+^ neurons and Npas1^+^ neurons are the two principal neuron classes in the GPe. A logistic regression with PV^+^ neurons ad Npas1^+^ neurons as the dependent variable and spontaneous rate and CV_ISI_ as the independent variables showed significant associations for both with neuron type—estimated odds ratio of 1.26 for spontaneous rate, *P* = 1.52 x 10^-7^, indicating a higher value in PV^+^ neurons; estimated odds ratio of 0.08 for CV_ISI_, *P* = 0.002, indicating a higher value in Npas1^+^ neurons. As logistic regression indicates a strong association between spontaneous activity level and neuron type, bootstrap analysis was applied to estimate the differences between genetically-defined neuron subtypes and these two principal neuron classes (Figure 10d). In general, 9 of the 10 characteristics considered - all except for AP height - show differences among cell types based on the Kruskal-Wallis test (Table 6), even after the Bonferroni multiple testing correction corresponding to a significance level of 0.05/10. It is notable that Npas1^+^-Lhx6^+^ neurons and Lhx6^+^_bright_ neurons have spontaneous activity that is statistically non-significant from Npas1^+^ neurons (*P* = 0.292 and 0.440, respectively). Furthermore, Dbx1^+^-PV^+^ neurons and Lhx6^+^_dim_ neurons exhibited spontaneous activity consistent with canonical PV^+^ neuron (*P* = 0.244 and 0.452, respectively). Pairwise comparisons of the electrophysiological characteristics across molecularly-defined GPe neuron subtypes are tabulated in Table 6.

As multiple electrophysiological characteristics were obtained, we asked whether a combination of these quantitative features can statistically define GPe neurons. To determine at the single-cell level whether genetically-defined GPe neuron subtypes form separate or overlapping clusters, we applied a principal component analysis to the electrophysiological dataset (Figure 11). Neurons with incomplete data were excluded from the analysis. As expected, and as illustrated by the dendrogram, neurons defined by PV and Npas1 transgenic lines were completely separated. Accordingly, they exhibited unique electrophysiological signatures as illustrated by the heatmap (Figure 11a). This analysis established the correspondence of electrophysiological classes with predefined markers, i.e. PV and Npas1. Moreover, different genetically-defined GPe neurons that were targeted in this study have electrophysiological properties that can be broadly categorized into either PV^+^ neuron and Npas1^+^ neuron classes. This analysis reaffirmed our previous assertion regarding the utility of these transgenic lines and markers (Hernandez et al., 2015). Though electrophysiological differences exist between Npas1^+^ neuron subclasses, hierarchical clustering of electrophysiological properties was insufficient to define GPe neuron subclasses.

**Figure 11.**
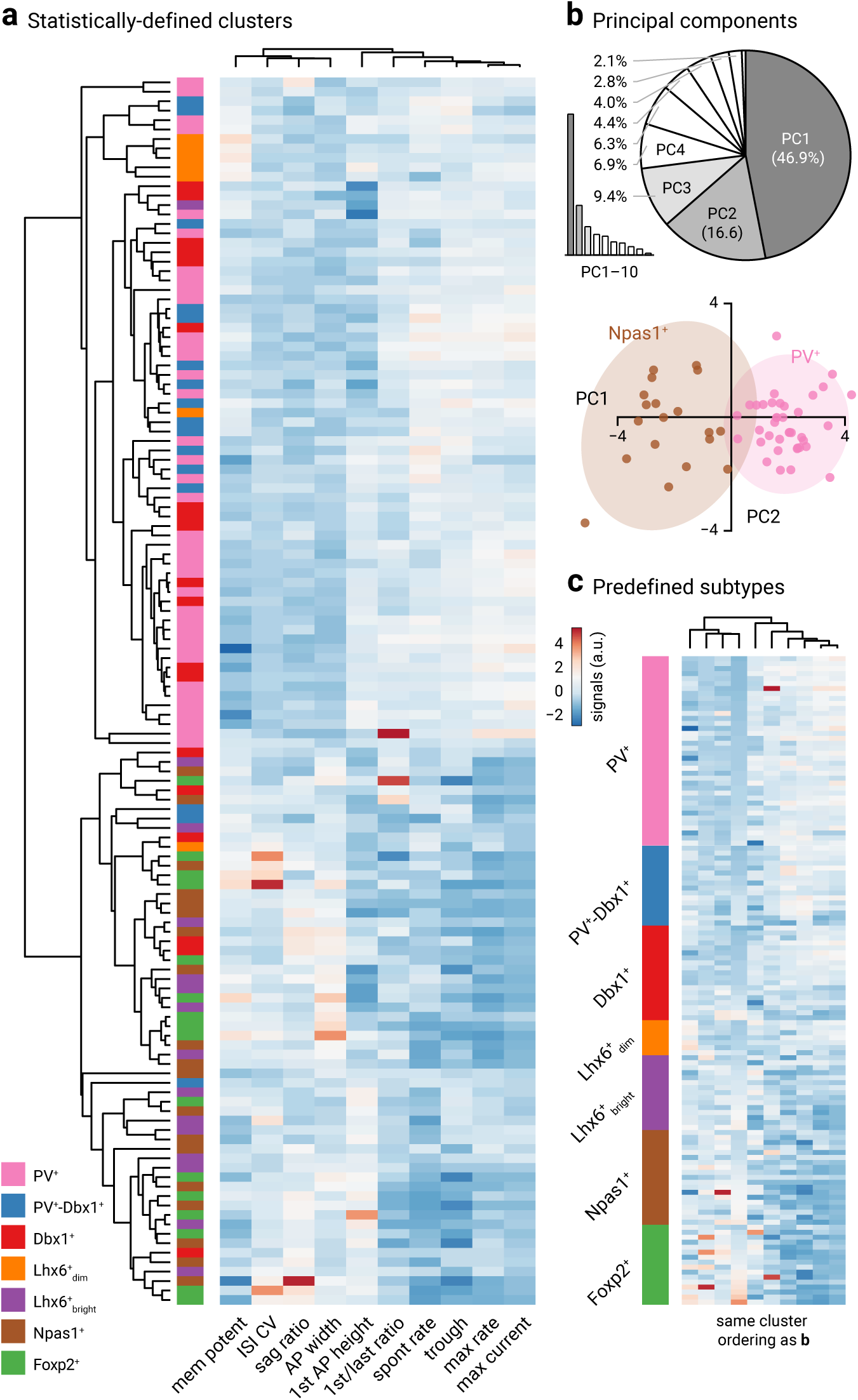
Electrophysiological multivariate analysis of GPe neurons. **a.** Heatmap representation of electrical signatures of genetically-identified GPe neuron subtypes. Dendrograms show the order and distances neuron clusters and their electrical characteristics. A total of 130 neurons (*n* = PV^+^: 38, Npas1^+^: 19, Dbx1^+^: 19, Foxp2^+^: 16, PV^+^-Dbx1^+^: 16, Lhx6^+^_bright_: 16, Lhx6^+^_dim_: 7) were included in this analysis. Neurons with incomplete data were excluded from the analysis. **b**. Top, scree plot (left) and pie chart (right) showing that the first three principal components (gray) capture 72.9% of the total variability in the data. Bottom, principal component 1 and 2 account for 46.9% and 16.6% of the total variance, respectively. Prediction ellipses for PV^+^ neurons (pink) and Npas1^+^ neurons (brown) are shown. With probability 0.95, a new observation from the same group will fall inside the ellipse. **c**. Same dataset as **a**. Data are sorted by genetically-identified neuron subtypes. Ordering of the clustering is the same as **a**.

Principal component analysis suggests that electrophysiological measurements covary with each other across GPe neurons; to gain insights into the relationships between these variables, we constructed a correlation matrix (Figure 12a, Table 7). The correlationship structure of these data provide the basis for the network analysis as shown in Figure 12b. This analysis shows good congruence with that from principal component analysis; importantly, they converge onto the idea that CV_ISI_, max current, and max rate are key defining features for GPe neurons (Figure 12c).

**Figure 12.**
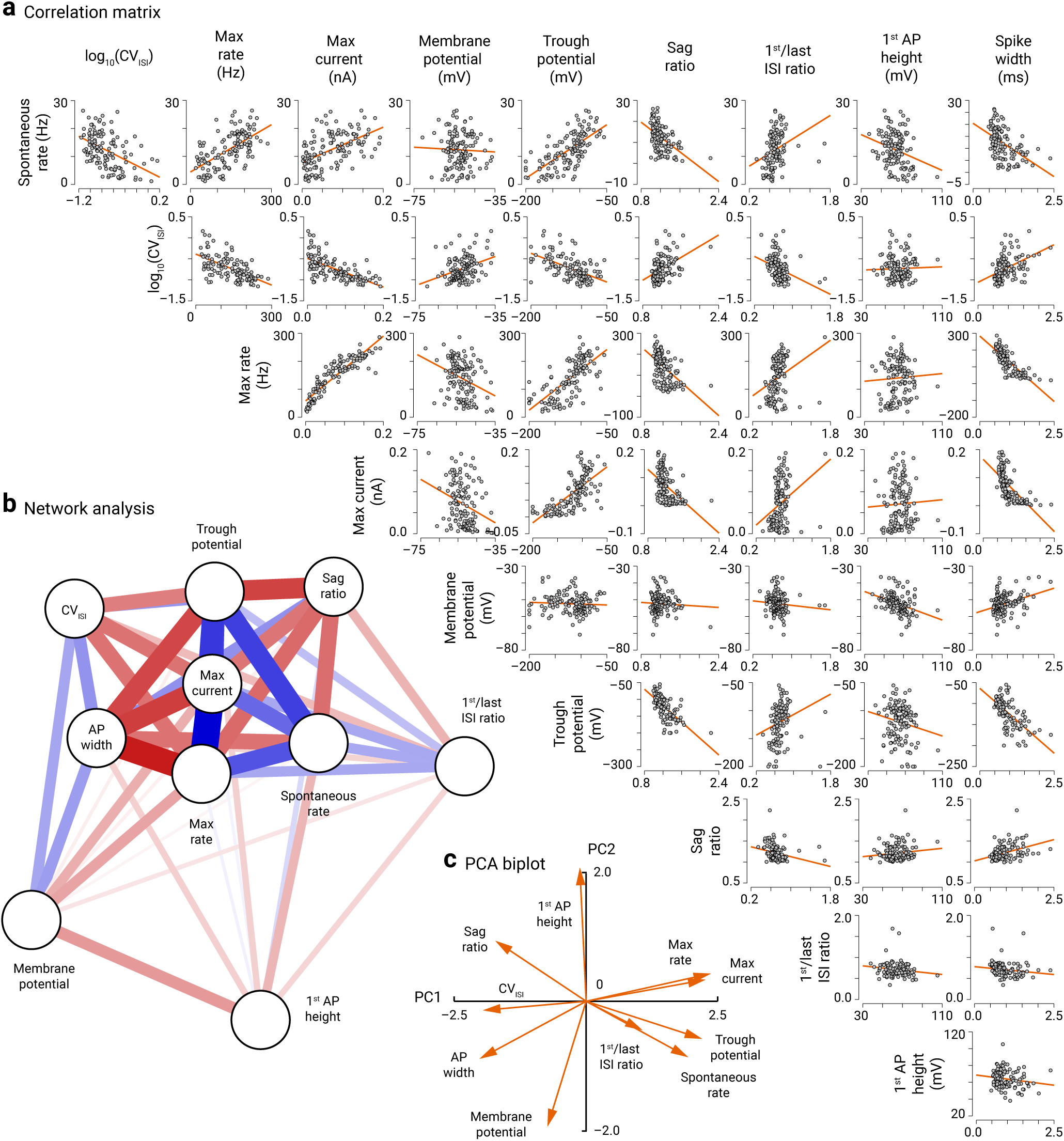
Correlation matrix and network analysis of electrophysiological measurements. **a.** A matrix table describing the pairwise correlation between all electrophysiological measurements. A total of 130 neurons (*n* = PV^+^: 38, Npas1^+^: 19, Dbx1^+^: 19, Foxp2^+^: 16, PV^+^-Dbx1^+^: 16, Lhx6^+^_bright_: 16, Lhx6^+^_dim_: 7) were included in this analysis. Orange lines represent the linear regression fits to the data in each plot. See Table 7 for Spearman *ρ* and *P* values. **b**. The strength of connections (aka edges) between nodes (white circles) are indicated by color and distance. Blue edges indicate positive relationships; red edges indicate negative relationships. The weights, lengths, and color saturation of lines indicate the strength of an edge. **c**. Principal component analysis biplot of electrophysiological measurements from recorded GPe neurons. The arrows (aka vector) represent eigenvectors showing the correlation of measurements to each other; measurements with a small angle between their vectors are strongly positively correlated, measurements with angles at 180° are expected to be strongly negatively correlated, and measurements perpendicular to each other (angles of 90 or 270°) are not correlated to each other. The vectors that point horizontally to the right are highly correlated with PC1 and the ones that point vertically upwards are highly correlated with PC2. As PC1 explains the majority of the variation (see Figure 11b) than PC2, the vertical arrows indicate measurements that are less defining. The length of a vector in the ordination plot reflects its contribution to the ordination. For simplicity, samples are not plotted.

**Table 7.** Correlation matrix of electrophysiological measurements

|  |  | Spearman correlations* |  |  |  |  |  |  |  |  |  |
| --- | --- | --- | --- | --- | --- | --- | --- | --- | --- | --- | --- |
|  |  | Spontaneous rate (Hz) | CV<sub>ISI</sub> | Max rate (Hz) | Max current (pA) | Membrane potential (mV) | Trough potential (mV) | Sag ratio | First/last ISI ratio | AP height (mV) | AP width (ms) |
| Spontaneous rate (Hz) | Spearman's $\rho$ | — | | | | | | | | | |
|  | *P* value | — |  |  |  |  |  |  |  |  |  |
| CV<sub>ISI</sub> | Spearman's $\rho$ | −0.476 | — | | | | | | | | |
|  | *P* value | < 0.001 | — |  |  |  |  |  |  |  |  |
| Max rate (Hz) | Spearman's $\rho$ | 0.621 | −0.619 | — | | | | | | | |
|  | *P* value | < 0.001 | < 0.001 | — |  |  |  |  |  |  |  |
| Max current (pA) | Spearman's $\rho$ | 0.558 | −0.688 | 0.934 | — | | | | | | |
|  | *P* value | < 0.001 | < 0.001 | < 0.001 | — |  |  |  |  |  |  |
| Membrane potential (mV) | Spearman's $\rho$ | −0.016 | 0.339 | −0.324 | −0.311 | — | | | | | |
|  | *P* value | 0.853 | < 0.001 | < 0.001 | < 0.001 | — |  |  |  |  |  |
| Trough potential (mV) | Spearman's $\rho$ | 0.717 | −0.576 | 0.741 | 0.763 | −0.106 | — | | | | |
|  | *P* value | < 0.001 | < 0.001 | < 0.001 | < 0.001 | 0.229 | — |  |  |  |  |
| Sag ratio | Spearman's $\rho$ | −0.54 | 0.453 | −0.558 | −0.546 | 0.095 | −0.709 | — | | | |
|  | *P* value | < 0.001 | < 0.001 | < .001 | < 0.001 | 0.284 | < 0.001 | — |  |  |  |
| First/last ISI ratio | Spearman's $\rho$ | 0.445 | −0.442 | 0.412 | 0.383 | −0.124 | 0.466 | −0.392 | — | | |
|  | *P* value | < 0.001 | < 0.001 | < 0.001 | < 0.001 | 0.161 | < 0.001 | < 0.001 | — |  |  |
| AP height (mV) | Spearman's $\rho$ | −0.225 | 0.029 | 0.062 | 0.115 | −0.441 | −0.136 | 0.056 | −0.162 | — | |
|  | *P* value | 0.01 | 0.744 | 0.485 | 0.192 | < 0.001 | 0.122 | 0.526 | 0.066 | — |  |
| AP width (ms) | Spearman's $\rho$ | −0.539 | 0.517 | −0.856 | −0.797 | 0.299 | −0.664 | 0.434 | −0.361 | −0.154 | — |
|  | *P* value | < 0.001 | < 0.001 | < .001 | < 0.001 | < 0.001 | < 0.001 | < 0.001 | < 0.001 | 0.081 | — |

\*Two-tailed *P* values are listed. Abbreviations: AP = action potential, CV = coefficient of variation; ISI = interspike interval, MAD = median absolute deviation.

## Discussion

In this study, we generated a more comprehensive landscape of GPe neuron composition. Specifically, we provided a more complete investigation of the Lhx6^+^ and Dbx1^+^ populations, along with novel insight into the properties of neurons arising from the Sox6 lineage. We were able to molecularly define the entirety of Lhx6^+^ neurons (see Table 3), thus resolving one of the debates in the field. Overall, our data further support the idea that PV^+^ neurons and Npas1^+^ neurons are two principal neuron classes. Our current study further illustrates the complexity of GPe neurons in adult mice and infers the existence of new neuron subclasses within Npas1^+^ neuron classes. Based on the molecular, anatomical, and intrinsic properties, our results support the idea that Npas1^+^-Nkx2.1^+^ neurons are a distinct GPe neuron subclass (Figure 13).

**Figure 13.**
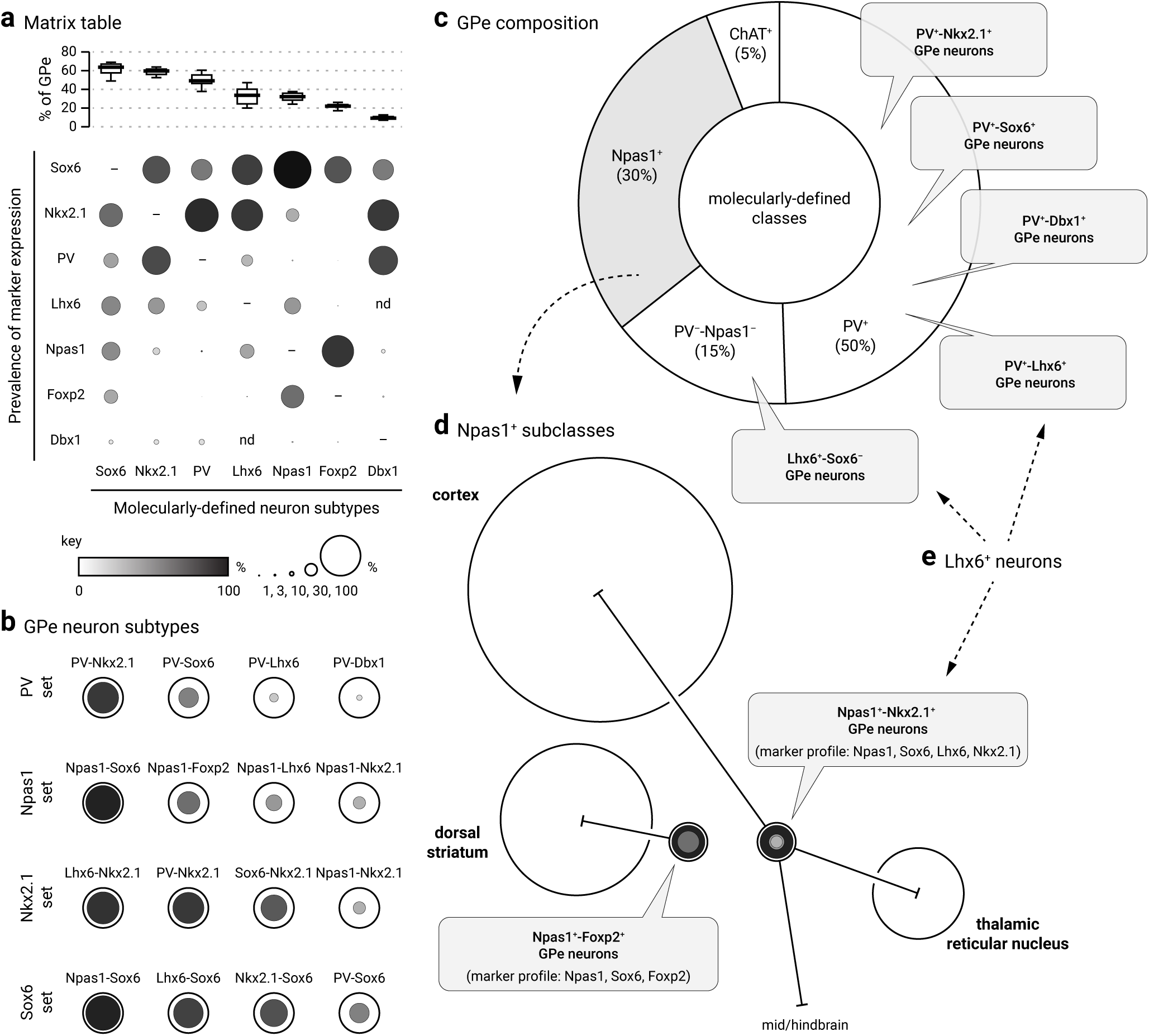
Diagrams summarizing the marker expression profile and classification scheme derived from the current study. **a.** The data from Table 3 is graphically represented to convey the co-expression of markers (vertical axis) within each molecularly-defined neuron subtype (horizontal axis). As visualized at the top of the matrix table, medians are presented as thick horizontal lines (see Figure 1) and data are sorted according to the abundance of neuron subtypes within the GPe. Both the size and grayscale intensity of each circle represents the prevalence of expression within specific GPe neuron subtypes. Circles along each column do not add up to 100% as there are overlapping markers expression with each neuron subtype. “–” denotes not shown; “nd” denotes not determined. For example, Foxp2 (6^th^ row) is selectively expressed in Npas1 neurons and a subset of Sox6 neurons; it is absent in Nkx2.1^+^ and PV^+^ neurons. Within the Foxp2^+^ population (6^th^ column), Sox6 and Npas1 are the only markers expressed. The high prevalence of Npas1 and Sox6 within the Foxp2^+^ neuron subtype (6^th^ column) is demonstrated by larger and darker circles. Note that because Foxp2 and Nkx2.1 are non-overlapping, there is no circle for Nkx2.1. Accordingly, Npas1^+^-Foxp2^+^ subclass and Npas1^+^-Nkx2.1^+^ subclass represent 57% and 32% of the Npas1^+^ population, respectively. As a comparison, Sox6 (1^st^ row) is expressed across all identified neuron types. Sox6 expression (1^st^ row) was observed in a major fraction of each population, especially Npas1^+^ neurons. The Sox6 neuron subtype (1^st^ column) expresses a broader range of markers. Lastly, neurons from the Dbx1 lineage (7^th^ row) are heterogeneous and overall contribute to only a small fraction of each molecularly-defined neuron subtype. **b.** A summary of the GPe neuron classification based on the expression profile of different molecular markers, i.e. data from **a**. **c**. A pie chart summarizing the neuronal composition of the mouse GPe. The area of each sector represents the approximate size of each neuron class. PV^+^ neurons (which constitute 50% of the GPe) are heterogeneous. Nkx2.1, Sox6, Lhx6, and Dbx1 are co-expressed in PV^+^ neurons to a varying extent. How they intersect with each other remains to be determined. Npas1^+^ neurons (gray) are 30% of the GPe; they can be subdivided into two subclasses (see **d**). ChAT^+^ neurons are ∼5% of the total GPe neuron population and show no overlap with other known classes of GPe neurons. **d**. Two *bona fide* subclasses of Npas1^+^ neurons (Npas1^+^-Foxp2^+^ and Npas1^+^-Nkx2.1^+^) are identified in the mouse GPe. They differ in their molecular marker expression, axonal projections, and electrophysiological properties. While Npas1^+^-Foxp2^+^ neurons project to the dorsal striatum, Npas1^+^-Nkx2.1^+^ neurons project to the cortex, thalamus, and mid/hindbrain areas. Size of the target areas (circles) is an artistic rendering based on the volume of those areas, but not the axonal density, synaptic strength, or contacts formed by Npas1^+^ neuron subclasses. **e**. Lhx6^+^ neurons are highlighted. Both PV^+^-Lhx6^+^ neurons and Npas1^+^-Lhx6^+^ neurons coexpress Sox6. Lhx6^+^-Sox6^−^ neurons are a subset of PV^−^-Npas1^−^ neurons.

### Classification of GPe neurons

Heterogeneity in the phenotype of GPe neurons was noted in the early 70’s (DeLong, 1971; Fox et al., 1974). The descriptions of molecularly-defined GPe neuron subtypes were not established until less than a decade ago (Flandin et al., 2010; Nobrega-Pereira et al., 2010). Our group has extended previous findings that PV^+^ neurons and Npas1^+^ neurons represent the two neuron classes in the adult GPe. Specifically, these two neuron classes are distinct across multiple modalities, including axonal patterns, electrophysiological properties, and alterations in disease states (Glajch et al., 2016; Hegeman et al., 2016; Hernandez et al., 2015). Our unpublished observations continue to support this notion; in particular, the synaptic inputs to PV^+^ neurons and Npas1^+^ neurons are distinct (Pamucku, Cui, Berceau, and Chan). On the other hand, other groups have adopted different, though not mutually exclusive, classification schemes.

Within the Npas1^+^ class, Foxp2^+^ neurons (Npas1^+^-Foxp2^+^, aka arkypallidal neurons) represent the first unique GPe neuron subclass to be described. This idea is supported by compelling data showing their distinct features, such as developmental origin, electrophysiological, anatomical, and molecular profiles (Dodson et al., 2015). Based on relative spike timing across basal ganglia nuclei in an operant behavioral task, it has been proposed that Npas1^+^-Foxp2^+^ neurons are important for motor suppression (Mallet et al., 2016). Meanwhile, the makeup of the remaining neurons in the GPe has been elusive. They are commonly referred to as prototypic neurons. We and others noted that prototypic neurons are not simply a single class of neurons (Abdi et al., 2015; Abrahao and Lovinger, 2018; Dodson et al., 2015; Flandin et al., 2010; Hernandez et al., 2015; Hunt et al., 2018; Mastro et al., 2014; Nobrega-Pereira et al., 2010; Oh et al., 2017; Saunders et al., 2018; Saunders et al., 2015). Instead, this group encompasses a heterogeneous population of neurons, and their properties have not been fully described. Our incomplete understanding of these neurons has prevented us from appreciating how individual neuron subclasses are precisely involved in motor function and dysfunction. Here, we found that Npas1^+^-Foxp2^+^ neurons are distinct from Npas1^+^-Nkx2.1^+^ neurons. Accordingly, both of them should be regarded as *bona fide* subclasses.

The examination of the molecular profiles of GPe neurons hinted at the diversity of PV^+^ neurons. Yet, we do not have sufficient data to argue if they fall into different neuron subclasses. Based on enhancer/transgenic mice, Silberberg and colleagues (Silberberg et al., 2016) suggest the existence of two pools of PV^+^ neurons that are produced with different temporal patterns and occupy slightly different, but otherwise largely overlapping spatial domains within the GPe. Consistent with this observation, more recent single-cell transcriptomic analysis confirms the existence of four major neuron types in the GPe, including two PV^+^ neuron clusters, in addition to two distinct Npas1^+^ neuron clusters (Saunders et al., 2018). It is paramount to determine whether they constitute distinct functional subclasses. These efforts will, in turn, give meaning to otherwise arbitrary classifications.

### Toward a more complete description of the GPe

We hypothesized the existence of a unique population of Lhx6^+^ neurons (i.e., Lhx6^+^-PV^−^-Npas1^−^) that accounts for at least 15% of GPe neurons. This idea is supported by our data along with data from others (Dodson et al., 2015; Hegeman et al., 2016; Hernandez et al., 2015; Mastro et al., 2014). In this study, we first examined the expression of Nkx2.1, Lhx6, and Sox6 among GPe neurons, as these transcription factors work in concert to dictate cell fate during development. As Sox6 is described as MGE-enriched, one would expect its expression in all Lhx6^+^ neurons. The examination of Sox6 expression unequivocally confirmed the existence of this unique Lhx6^+^ population— however, to our surprise, this unique Lhx6^+^ population is Sox6^−^. Importantly, our results resolve some of the discrepancies related to Lhx6^+^ neurons and identify the PV^−^ and Npas1^−^ neurons described in our previous study (Dodson et al., 2015; Hegeman et al., 2016; Hernandez et al., 2015; Mastro et al., 2014).

While Dbx1^+^ neurons that originate from the PoA are known to populate the GPe (Gelman et al., 2009; Nobrega-Pereira et al., 2010), their properties were not fully-determined. We examined whether Dbx1^+^ neurons correspond to the Lhx6^+^-Sox6^−^ population. Contrary to our hypothesis, our data argue that Dbx1^+^ neurons do not correspond to this population of neurons. Instead, we found the Dbx1^+^ population contains neurons that express, to varying degrees, all established GPe neuron markers (Figure 2, Table 3). This is consistent with the literature that PoA progenitors give rise to a small, but diverse, contingent set of neurons (Gelman et al., 2011; Gelman et al., 2009). In particular, most Dbx1^+^ neurons are Sox6^+^, and they primarily express PV^+^ or Npas1^+^. In hindsight, these results were not completely unexpected. While the embryonic PoA is similar to the MGE in that it expresses the transcription factor Nkx2.1, many PoA cells do not express Lhx6 (Flames et al., 2007). It has been shown that a subset of LGE (lateral ganglionic eminence) progenitors express Lhx6 (Liodis et al., 2007). Importantly, the LGE is known to generate GABAergic neurons that populate the cortex (Anderson et al., 2001; de Carlos et al., 1996; Jimenez et al., 2002; Tamamaki et al., 1997). Therefore, it is possible that the Lhx6^+^-Sox6^−^ GPe population arises from the LGE. One limitation with using the Lhx6-GFP mouse, which is a BAC transgenic line, is the concern that GFP is expressed in ectopic neurons. However, given the near complete overlap with Nkx2.1, we are confident that its expression is accurate within the GPe. We await new tools that give us unique access to the Lhx6^+^-Sox6^−^population. It is interesting to see that PV^+^-Dbx1^+^ neurons display a phenotype that is shared with the general PV^+^ population, which originate primarily from the MGE (Flandin et al., 2010; Nobrega-Pereira et al., 2010). Although the extent of the overlap remains to be fully established, our findings are in line with what was shown previously that neurons that arise from spatially-distinct ontogenic domains can converge onto a single neuron class (Chittajallu et al., 2013). However, cellular phenotypes, such as transcriptomic program and electrophysiological profiles, can be shaped by neuromodulatory signals. It is intriguing to hypothesize that brain states, imposed by various neuromodulatory signals, may have differential impacts on these neuron subtypes as a result of distinct receptor profiles. High-throughput single-cell transcriptomic analysis has become an extremely powerful tool for cell classification. However, as both Lhx6^+^-Sox6^−^ and Dbx1^+^ neurons are sparse in the GPe, they are likely underrepresented in previous single-cell transcriptomic studies (Saunders et al., 2018; Zeisel et al., 2018). Our current study has thus provided important insight into these low abundance neurons.

### Significance of the cortico-pallido-cortical loop

Though our connectome survey did not reveal either Lhx6^+^-Sox6^−^ neurons or Dbx1^+^ neurons to be cortically projecting, it pinpointed Npas1^+^-Nkx2.1^+^ neurons as key constituents of the cortico-pallido-cortical loop. This finding can have far-reaching implications in motor planning, motor learning, and decision making (Barthas and Kwan, 2017; Hanks and Summerfield, 2017; Ito and Doya, 2011; Papale and Hooks, 2018; Svoboda and Li, 2018). We noted that the GPe innervation is not a consistent feature of all PT neurons (Cowan and Wilson, 1994; Kita and Kita, 2012; Shepherd, 2014; Shibata et al., 2018), this is likely related to the rich diversity of cortical neuron subtypes (Gouwens et al., 2019).

Importantly, recent data have shown that the electrophysiological activity of primate PT neurons is altered in the parkinsonian state (Pasquereau and Turner, 2011). Our results provide insight into a potential cellular mechanism that underlies the effectiveness of deep brain stimulation (DBS), which involves the implantation of a frequency stimulation electrode in the STN. DBS has emerged as a crucial adjunct to manage the motor symptoms of PD (Wichmann et al., 2018; Wichmann and Delong, 2006). The present results suggest that activation of cortical axons during DBS may activate brain sites innervated by the multi-projectional PT-type cortical neurons. Activation of these brain regions, including the GPe, may contribute to the alleviation of motor symptoms with DBS.

### Concluding remarks

In this study, we have performed a thorough characterization of GPe neuron populations according to their expression of genetic markers, electrophysiological properties, and anatomical projections. In addition, we have attempted to characterize and employ various novel driver and reporter lines. We hope that our findings will facilitate cross-laboratory utilization of standard tools to study GPe neuron types. The identification of GPe neuron populations should allow experiments to be conducted on the same neuron type across subjects and laboratories. Examining the same (i.e., homologous) neuron population across species facilitates comparative studies; commonalities and differences in phenotype could then be linked to behavior. As we have completed cataloging major neuron types within the GPe, our next goal is to use intersectional tools to define constituent neuron subclasses and their functions. We have used similar strategies in this study and also recently in Poulin et al. (Poulin et al., 2018). The generation and identification of additional Flp driver lines will likely be helpful for the interrogation of GPe neuron diversity and function. Ultimately, our goal is to identify single recombinase driver lines that efficiently capture functional neuron subclasses.

## Acknowledgments

This work was supported by NIH R01 NS069777 (CSC), P50 NS047085 (CSC), R01 MH112768 (CSC), R01 NS097901 (CSC), R01 MH109466 (CSC), R01 NS088528 (CSC), R01 NS096240 (RA), R01 MH110556 (RA), P50 DA044121 (RA), R01 MH107742 (BL), R01 MH108594 (BL), U01 MH114829 (BL), R01 MH116176 (YK), R01 MH112768 (NJJ), R56 MH114032 (NJJ), R21 AA026022 (NJJ), R01 NS103993 (BMH), ZIA MH002497 (CRG), T32 NS041234 (HSX), F32 NS098793 (HSX), T32 AG020506 (AP), NARSAD Young Investigator Award (BMH), HHMI-PF Medical Research Fellowship (ZA), AΩA Student Research Fellowship (ZA) and Northwestern University Weinberg Summer Research Grant (PHW). We thank Xixun Du, Yu Zhang, Vishnu Rangachari, Kris Shah, Elizabeth Augustine, and Daniel Hegeman for their assistance on the project, Alexandria Granados, Morgan Marshall, and Nicole Curtis, for colony management and technical support, Dr. Alicia Gumez-Gamboa for providing Emx1-Cre mice, Dr. Jeffrey Savas for providing VGluT1 antibody, and Northwestern University Transgenic and Targeted Mutagenesis Laboratory for providing EIIa-Cre and CAG-Flp breeders.

